# *Theobroma cacao* improves bone growth by modulating defective ciliogenesis in a mouse model of achondroplasia

**DOI:** 10.1101/2021.02.18.431801

**Authors:** L. Martin, N. Kaci, C. Benoist-Lasselin, M. Mondoloni, S. Decaudaveine, V. Estibals, M. Cornille, L. Loisay, J. Flipo, B. Demuynck, M. de la Luz Cádiz-Gurrea, F. Barbault, S. Fernández-Arroyo, L. Schibler, A. Segura-Carretero, E. Dambroise, L. Legeai-Mallet

**Affiliations:** Université de Paris, Imagine Institute, Laboratory of Molecular and Physiopathological Bases of Osteochondrodysplasia. INSERM UMR 1163, F-75015, Paris, France; Inovarion, F-75005 Paris, France; Department of Analytical Chemistry, University of Granada, S-18100 Granada, Spain; Research and Development of Functional Food Centre (CIDAF), S-18100 Granada, Spain; Université de Paris, ITODYS, CNRS, UMR 7086, 15 rue J-A de Baïf, F-75013 Paris, France; Biomedical Research Unit. Medicine and Surgery Department, Rovira i Virgili University, S-43002 Tarragona, Spain; ALLICE, Maison Nationale des Eleveurs, F-75012 Paris, France

## Abstract

A gain-of-function mutation in the fibroblast growth factor receptor 3 gene (*FGFR3*) results in achondroplasia (ACH), the most frequent form of dwarfism. The constitutive activation of FGFR3 impaired bone formation and elongation and many signaling transduction pathways. Identification of new and relevant compounds targeting the FGFR3 signaling pathway is of broad importance for the treatment of ACH. Natural plant compounds are the prime sources of drug candidates. Here, we found that the phenol compound (-)-epicatechin isolated from *Theobroma cacao* effectively inhibits FGFR3’s downstream signaling pathways. Transcriptomic analysis in *Fgfr3* mouse model showed that ciliary mRNA expression was modified and influenced significantly by the Indian hedgehog and PKA pathways. (-)-Epicatechin is able to rescue impairments in the expression of these mRNA that control both the structural organization of the primary cilium and ciliogenesis-related genes. In femurs isolated from a mouse model (*Fgfr3*^*Y367C/*+^) of ACH, we showed that (-)-epicatechin countered the bone growth impairment during 6 days of *ex vivo* cultures. We confirmed *in vivo* that daily subcutaneous injections of (-)-epicatechin in *Fgfr3*^*Y367C/*+^ mice increased bone elongation and rescued the primary cilium defect observed in chondrocytes. This modification of the primary cilia promoted the typical columnar arrangement of flat proliferative chondrocytes and thus enhanced bone elongation. The results of the present proof-of-principle study illustrated (-)-epicatechin’s ability may facilitate the development of (-)-epicatechin as a treatment for patients with ACH.

## Introduction

The most common form of dwarfism, achondroplasia (ACH), is caused by a gain-of-function mutation in the *FGFR3* gene coding for fibroblast growth factor receptor 3^1^. FGFR3 gain-of-function mutations are also associated with hypochondroplasia, a mild dwarfism and thanatophoric dysplasia (TD) a severe and lethal dwarfism^2^.

The defective FGFR3 signal transduction impairs intracellular downstream signaling pathways including extracellular signal-regulated kinases-1 and 2 (Erk1-2) and p38, phosphoinositide 3-kinase/Protein kinase B (PI3K/AKT), phospholipase Cγ (PLCγ) and Signal Transducer and Activators of Transcription (STATs) and thus affects chondrocyte proliferation, differentiation and bone elongation^2^. FGFR3 plays a significant role in the growth plate development, acting to inhibit both the rate of chondrocyte proliferation and differentiation and interacts with Indian hedegehog (Ihh) signaling pathway to control chondrocyte formation^3^. FGF signaling regulates the length of primary cilia in many tissues^4,5^. FGFR3 signaling interacts with Hedgehog and the serine/threonine kinase intestinal cell kinase (Ick), which is involved in ciliogenesis and participate in control in ciliary length^6,7^. In ACH, we demonstrated a defective primary cilium elongation in mouse and human chondrocytes^6–8^. The regular alignment of primary cilia is responsible of columnar-stacked chondrocytes in growth plate cartilage^9^. Primary cilium is primordial for the regulation of chondrocyte rotation, as demonstrated by the presence of defective primary cilium biosynthesis and/or function in many skeletal ciliopathies^10,11^.

In recent years, a significant body of work has focused on treatments for ACH. Non-surgical therapeutic strategies have been developed from insights gained from preclinical studies using *Ach* mouse models^2,12^. The therapeutic approaches are many and varied, and include (*i*) targeting FGF ligand (Recifercept) and aptamer (APT-F2P/RBM 007), (*ii*) targeting receptor FGFR (anti-FGFR3 antibody-B701/vofatamab), (*iii*) inhibiting the tyrosine kinase activity (BGJ398/Infigrantinib) or (*iv*) using C-type natriuretic peptide (CNP) (TransCon CNP, BMN111/Vosoritide) analog^13^ to antagonize the mitogen-associated protein kinase (MAPK) pathway. Vosoritide approach is currently furthest along the clinical development pathway^14,15^.

Identification of new and relevant compounds targeting the FGFR3 signaling pathway is of broad importance for the treatment of FGFR3-related chondrodysplasia. Natural plant compounds are the prime sources of drug candidates^16^. Plant polyphenols such as *Theobroma cacao* contains flavon-3-ols and polyphenols that have long been considered to have relevant biological activities in the treatment of a number of diseases and that are known to act in various ways on MAPK signaling pathways, *e*.*g*. by (*i*) reducing reactive oxygen species production and Erk1-2 and p38 phosphorylation in neurons^17^, (*ii*) inhibiting adipocyte differentiation through AMP activated protein kinase (AMPK) and Erk1-2 signaling pathways^18^, and (*iii*) modulating antioxidant enzyme activities through Erk1-2 and regulating glucose production through AMPK modulation in liver cells^19^.

In this study, we identified (-)-epicatechin from a *Theobroma cacao* extract as drug candidate. (-)-Epicatechin is able to (*i*) induce primary cilium elongation of chondrocytes and modified the ciliary genes expression that were differentially expressed in the growth plate cartilage of *Fgfr3*^*Y367C/*+^ mice (*ii*) decrease FGFR3’s downstream signaling pathways in cell and organ cultures and (*iii*) promote femur growth elongation in both femur explants and into live *Fgfr3*^*Y367C/*+^ mice. Our results highlighted this molecule’s specific action on the Ihh pathway related to primary cilia, and suggest that (-)-epicatechin could be developed as a means of correcting bone growth defects in patients with achondroplasia.

## Results

### *Theobroma cacao* extract restores the abnormal activation of FGFR3 pathway and primary cilium defect in mutant *Fgfr3* chondrocytes

To investigate the therapeutic efficacy of *Theobroma cacao* on the abnormal activation of FGFR3 pathway, we fractionated *Theobroma cacao* extract by combining solid-phase extraction with semi-preparative HPLC^16^ and collected a total of 11 fractions (Fraction 1 to 11) (Fig. 1a). The activity of the fractions was tested according to two relevant criteria considering as an hallmark of FGFR3-related disorders namely *i*) the ability to inbihit the MAPK pathway a downstream effector of the activated FGFR3 pathway^20^ *ii*) the ability to promote primary cilia elongation^6,8^.

**Figure 1.**
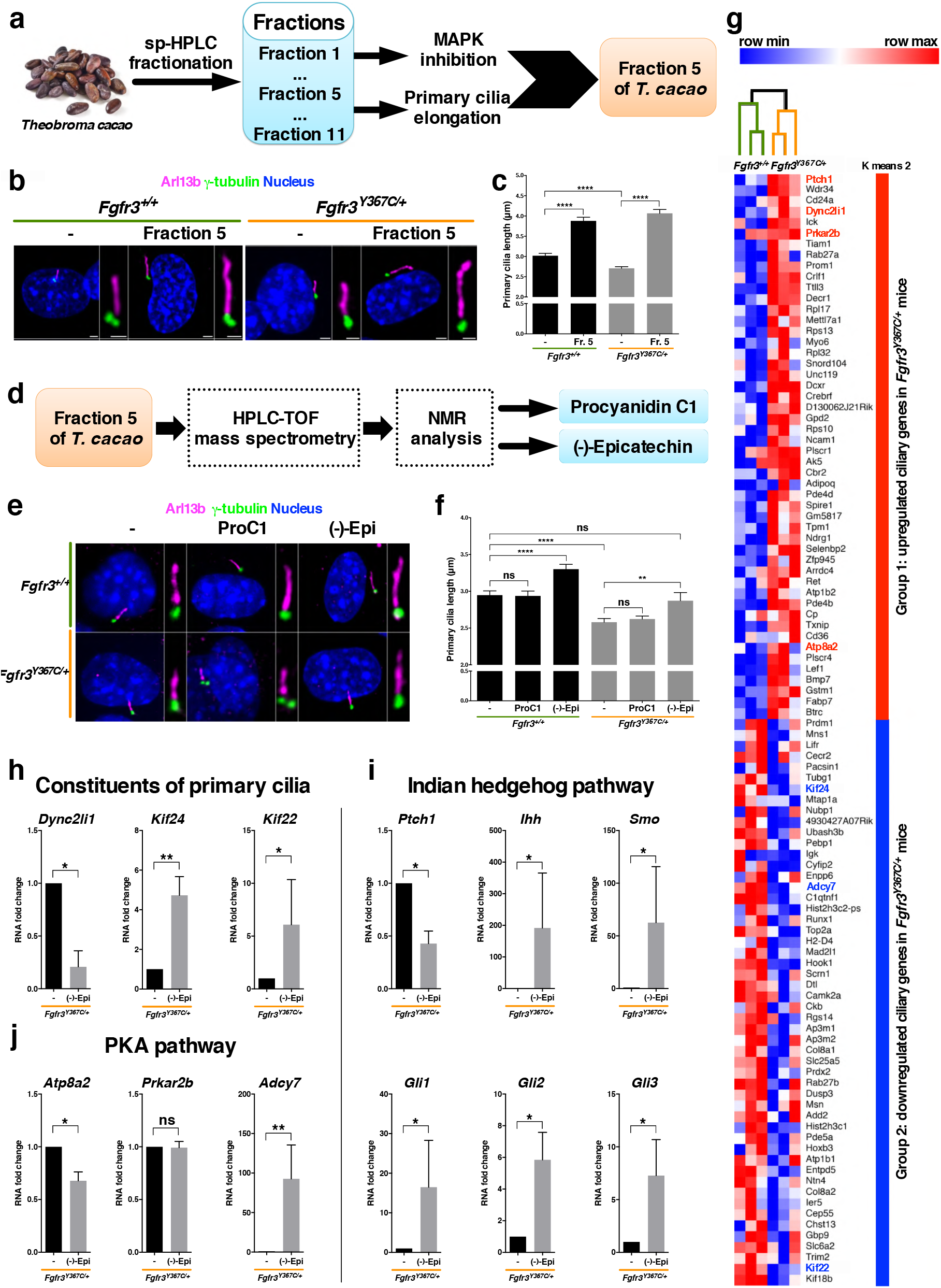
(-)-Epicatechin from fraction 5 of *Theobroma cacao* modifies both ciliary length and ciliary gene expression. **a**, Sequential representation of fractionation and selection of *Theobroma cacao*. **b**, Representative confocal microscopy image of primary cilium immunolabelled with Arl13b (magenta) for axoneme, γ-tubulin (green) for basal body, and DAPI (blue) for cell nuclei in E16.5 chondrocytes. **c**, Graphical representation of the length of the primary cilium in E16.5 *Fgfr3*^+*/*+^ (n=219 and n=162 for exposed cells) and *Fgfr3*^*Y367C/*+^ murine chondrocytes exposed (n=338) or not (n=443) to fraction 5 of *Theobroma cacao*. **d**, Sequential representation of identification of two major compounds (procyanidin C1 and (-)-epicatechin) of *Theobroma cacao* fraction 5. **e**, Representative confocal microscopy image of the E16.5 primary cilium immunolabeled with Arl13b (magenta), γ-tubulin (green), and DAPI (blue). **f**, Graphical representation of the length of the primary cilium in E16.5 *Fgfr3*^+*/*+^ not exposed (2.95 ± 0.06; n=300), exposed to procyanidin C1 (2.94 ± 0.07; n=232) or to (-)-epicatechin (3.30 ± 0.07; n=278) and *Fgfr3*^*Y367C/*+^ murine chondrocytes not exposed (2.58 ± 0.05; n=272), exposed to procyanidin C1 (2.62 ± 0.04; n=461) or to (-)-epicatechin (2.87 ± 0.11; n=113) **g**, Heat map from a comparative transcriptome analysis of upregulated (red) and downregulated (blue) genes of growth plate cartilage from 1-week-old *Fgfr3*^+*/*+^ and *Fgfr3*^*Y367C/*+^ mice. **h**, RNA expression fold-change for *Dync2li1, Kif24* and *Kif22* mRNA. **i**, RNA expression fold-change for *Ptch1, Ihh, Smo, Gli1, Gli2* and *Gli3* mRNA. **j**, RNA expression fold-change for *Atp8a2, Prkar2* and *Adcy7* mRNA. Data are quoted as the mean ± SEM. ns, not significant; *\*p<0*.*05*; *\*\*p<0*.*01*; *\*\*\*p<0*.*001*; *\*\*\*\*p<0*.*0001* in a two-tailed, unpaired t-test.

We found by western-blot that *Theobroma cacao* extract fraction 5 significantly lowered Erk1-2 phosphorylation in human ACH and TD chondrocytes (Supplementary Fig. 1a,b). Moreover, we found, by immunolabelling of chondrocytes isolated from *Fgfr3*^*Y367C/*+^mice exhibiting a dwarf phenotype^21^ that fraction 5 increased the initial primary cilium length defect in *Fgfr3*^*Y367C/*+^ chondrocytes treated (4.06 ± 0.10 µm), compared to non-treated *Fgfr3*^*Y367C/*+^ chondrocytes (2.71 ± 0.04 µm) (Fig. 1b,c).

### Two compounds procyanidin C1 and (-)-epicatechin are present in fraction 5

To identify the putative dual-effect compound, the compounds comprised in fraction 5 from *Theobroma cacao* were separated and purified by HPLC system coupled to a time-of-flight (TOF) mass spectrometer and analysed by NMR (Fig. 1d). The NMR analyses revealed the presence of two different compounds: procyanidin C1 (10-20%) and (-)-epicatechin (80-90%) (Fig. 1d and Supplementary Fig. 2a-e). To evaluate, the ability of both procyanidin C1 and (-)-epicatechin to act on primary cilium elongation, we measured the length of primary cilia in *in vitro Fgfr3*^*Y367C/*+^ and *Fgfr3*^+*/*+^ cultured mouse chondrocytes (Fig. 1e). Treatment with procyanidin C1 (with a range of concentrations, not showed) failed to increase primary cilium length in both *Fgfr3*^*Y367C/*+^ chondrocytes (2.62 ± 0.04 µm) and *Fgfr3*^+*/*+^ (2.94 ± 0.07 µm) compared to non-treated *Fgfr3*^*Y367C/*+^ (2.58 ± 0.05 µm) and non-treated *Fgfr3*^+*/*+^ chondrocytes (2.95 ± 0.06 µm), respectively (Fig. 1e,f). Interestingly, treatment with (-)-epicatechin rescued the primary cilium length defect in *Fgfr3*^*Y367C/*+^ chondrocytes; the mean length (2.87 ± 0.11 µm) was similar to that measured in non-treated *Fgfr3*^+*/*+^ chondrocytes (2.95 ± 0.06 µm) (Fig. 1e,f). Here, we conclude that (-)-epicatechin is the sole active compound present in *Theobroma cacao* fraction 5 extract showing a benefitial action on primary cilia lenght.

### Transcriptomic identification of primary cilium genes in cartilage

The mechanism by which FGF signaling controlling cilia in growth plate cartilage is not well understood. Here, in order to define relevant genes controlling the ciliogenesis in FGFR3-related dwarfism, we performed a comparative, longitudinal, transcriptomic analysis of RNA isolated from cartilage growth plate of *Fgfr3*^*Y367C/*+^mice *versus* control. After the identification of 3,336 modulated probe sets, 763 differentially expressed probe sets (612 genes) were found to be significantly modulated (absolute fold change > 1.5, and an adjusted *p* value ≤ 5% relative to overall changes in expression) and were fed into a functional analysis (Gene Expression Omnibus, database ID: GSE145821). Of these, 102 genes were shown to be related to primary cilia (Gene Expression Omnibus database ID: GSE145821, Gene Ontology annotation and/or registration in the Cildb primary cilia gene database)^22^. As shown in Fig. 1g, hierarchical clustering on expression levels led to the identification of two groups of upregulated (group 1) or downregulated (group 2) genes. In agreement with previous data, we found that Ick was upregulated^7^ as well as we pointed out relevant upregulating primary cilia genes such as Ptch1 (protein patched homolog 1), Dync2li1(coding for dynein 2 light intermediate chain 1, a regulator of primary cilium length), Prkar2b (Protein Kinase cAMP-dependent Type II Regulatory Subunit β); and Atp8a2 (ATPase Phospholipid Transporting 8A2). Interestingly, we observed the downregulation of Kif24 (kinesin family member 24), Adcy7 (Adenylate Cyclase 7) and Kif22 (kinesin family member 22). Among the selected genes upregulated or downregulated, we were able to distinguish three families of genes associated with *i*) cilium components, *ii*) the Hh (Hedgehog) pathway and *iii*) the protein kinase A (PKA) pathway (Fig. 1g). In an attempt to further understand the mechanisms underlying (-)-epicatechin’s effects on the primary cilium, using *in vitro* test we monitored gene expression in *Fgfr3*^*Y367C/*+^ chondrocytes treated or not with (-)-epicatechin (Fig. 1h-j). We first evaluated the genes *Dync2li1, Kif24* and *Kif22* involved in cilia formation and skeletal ciliopathies (Fig. 1h)^23–25^. *Dync2li1* is highly expressed in *Fgfr3*^*Y367C/*+^ chondrocytes^25^, and this expression was significantly reduced 4.8-fold by (-)-epicatechin treatment. The lower expression levels of *Kif24* and *Kif22* observed in *Fgfr3*^*Y367C/*+^ growth plate (Fig. 1g) were increased by (-)-epicatechin treatment of *Fgfr3*^*Y367C/*+^ chondrocytes by 4.7- and 6.1-fold, respectively (Fig. 1h). Secondly, we investigated the Hh pathway commonly dysregulated in ciliopathies^26,27^ (Fig. 1i). It is well established that primary cilia play a central role in Ihh signal transduction. Ihh binds to Ptch1 receptor to release Smo (*Smoothened)* signal transducer from Patched-dependent suppression. Smo stabilizes Gli (*Gli family zinc finger)* family members and promotes nuclear accumulation of Gli. The elevated mRNA expression of the *Patch1* in *Fgfr3*^*Y367C/*+^ chondrocytes (Fig. 1g) was notably reduced by 2.3-fold in (-)-epicatechin-treated chondrocytes (Fig. 1i). The downregulated expression of *Ihh* and successive downstream effectors of *Ptch1*: *Smo, Gli1, Gli2* (*Gli-Kruppel family member 2*) and *Gli3* (*Gli-Kruppel family member 3*) in *Fgfr3*^*Y367C/*+^ chondrocytes compared to controls, were strongly increased by (-)-epicatechin treatment (by 191.7-, 62.6-, 16.5-, 5.9- and 7.3-fold, respectively) (Fig. 1i). We investigated the PKA pathway as a regulator of Hh signaling^28^. Interestingly, the overexpression of *Atp8a2* in mutant chondrocytes was reduced 1.5-fold by (-)-epicatechin treatment (Fig. 1j). In contrast, overexpression of *Prkar2b* (another member of the PKA-dependent pathway) in *Fgfr3*^*Y367C/*+^ chondrocytes was not modified by the treatment (Fig. 1j). Furthermore, we observed that *Adcy7* expression was increased 92.7-fold by (-)-epicatechin treatment (Fig. 1j). Our results suggest that in FGFR3-related disorders, (-)-epicatechin is able to rescue impairments in the expression of (*i*) key genes that control both the structural organization of the primary cilium and (*ii*) ciliogenesis-related genes that act on the Ihh and PKA signaling pathways involved in the regulation of cartilage homeostasis.

### (-)-Epicatechin counteracts the abnormal activation of FGFR3 signaling pathway

Since (-)-epicatechin had shown beneficial effects on primary cilia formation in the *Fgfr3*^*Y367C/*+^ chondrocytes and in order to determine whether (-)-epicatechin acts directly on FGFR3 phosphorylation, we transiently transfected human embryonic kidney 293 (HEK293) cells with FGFR3 constructs bearing a gain-of-function mutation in the protein’s extracellular domain (TD/Y373C; Fig. 2a), transmembrane domain (ACH/G380R; Fig. 2b) or intracellular domain (TD/K650E; Fig. 2c). FGFR3 phosphorylations in mutants were not modified by (-)-epicatechin treatment (Fig. 2a-c). We next performed *in silico* analysis to define (-)-epicatechin’s ability to bind to FGFR3 kinase’s binding pocket. Visualization of a structural model showed clearly that (-)-epicatechin does not insert fully into the FGFR3’s ATP-binding pocket. To check the ability of (-)-epicatechin to bind the kinase target, a molecular dynamics simulation (duration: 50 ns) was undertaken for each (-)-epicatechin complex. The results of the hyperdynamics calculations (static state to dynamic state, Fig. 2d) revealed that (-)-epicatechin was unable to form a favorable complex with human FGFR3 and was gradually pushed away from the FGFR3 binding pocket (Fig. 2d, bottom panel and Supplementary Table 1). During the first 10 nanoseconds (Fig. 2e, left panel), (-)-epicatechin slowly broke its hydrogen bonds to Y557, E556 and A558 and became more exposed to the solvent (water). At the end of the “conventional” 50 ns molecular dynamics step, (-)-epicatechin was still at the surface of the FGFR3 protein but was too far from the kinase’s active site (Fig. 2e, last right panel) to act as inhibitor (for detailed animation, please see Supplementary Movie 1). In light of these results, we performed similar experiments with procyanidin C1, we observed that this large molecule did not interact with FGFR3 (Supplementary Fig. 3 and Supplementary Table 2).

**Figure 2.**
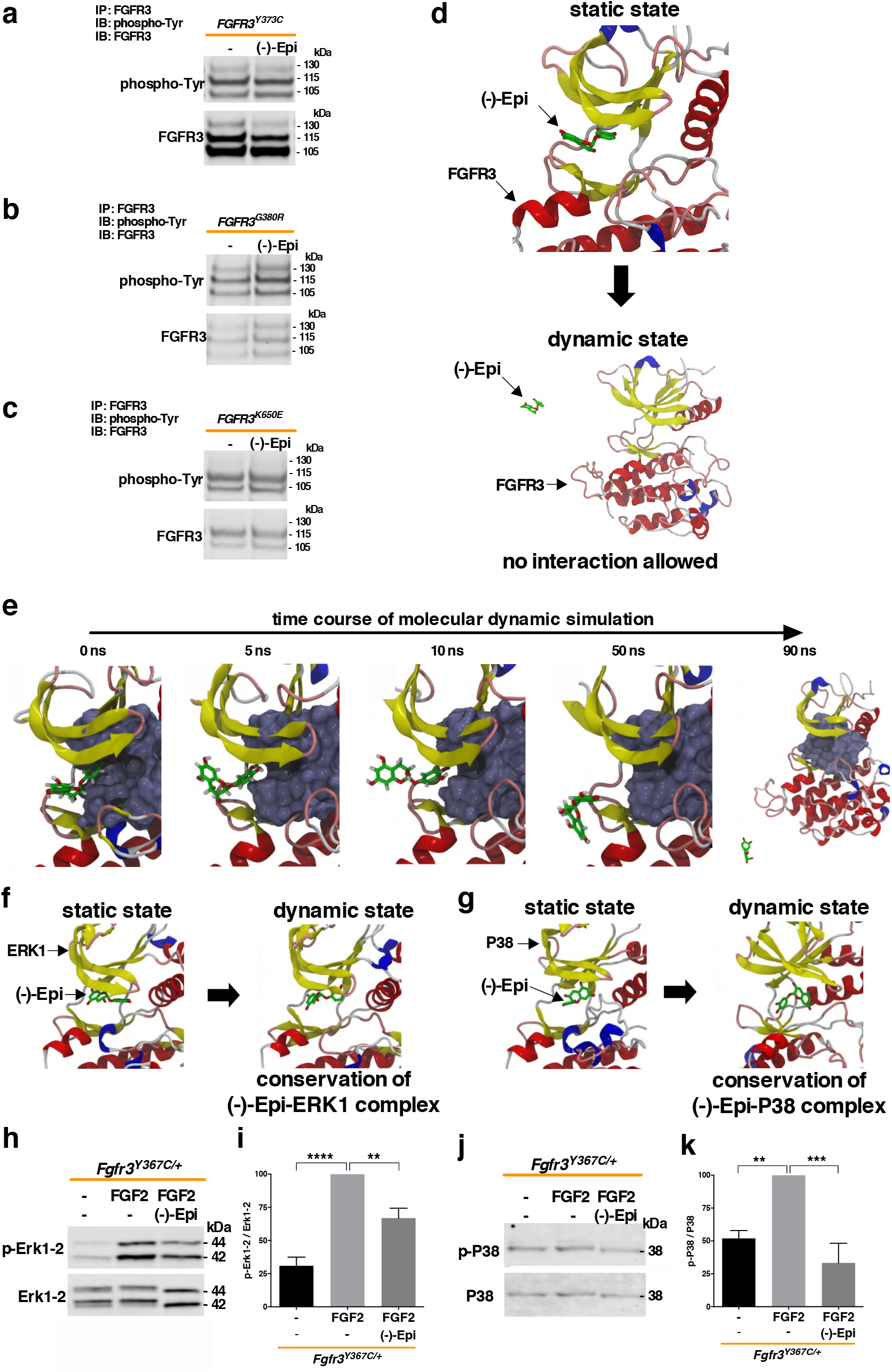
(-)-Epicatechin acts downstream of FGFR3 on cartilage homeostasis. **a, b, c**, Representative western-blots of tyrosine-phosphorylated FGFR3 (p-Tyr) expression from FGFR3 immunoprecipitation of HEK293 cells transfected with *FGFR3*^*Y373C*^ (**a**), *FGFR3*^*G380R*^ (**b**) and *FGFR3*^*K650E*^ (**c**) exposed or not to (-)-epicatechin. From (**d**) to (**g**): computational results for (-)-epicatechin’s binding to FGFR3. In all these pictures, the protein is displayed as ribbons (yellow: β-sheet, red: α-helix, pink: turn and white: random coil) and the ATP active site is presented as a molecular surface in ice blue. The (-)-epicatechin is displayed in CPK colors, with carbon in green. Structures on the top (**d**, top) correspond to the “FGFR3-(-)-epicatechin” complex after the molecular docking computations. The hydrogen bonds between interacting residues (presented with grey carbon atoms) are shown with dash lines. The structure on the bottom (**d**, bottom) was obtained after the molecular dynamics and hyperdynamics stages. **e**, Detailed illustrations of the dissociation of (-)-epicatechin from FGFR3 kinase. From left to right, snapshots were extracted after 0, 5, 10 and 50 ns of classical molecular dynamics and after 40 ns of the hyperdynamics stage (See the Supplemental Movie 1). **f, g**, Representative structures of (-)-epicatechin complexed with ERK1 (**f**), ERK2 (Supplementary Fig. 4) and P38 (**g**) kinases. The complexes obtained after the molecular docking computations (*i*.*e*. static states) are shown on the left, and those obtained after the molecular dynamics and hyperdynamics simulation stages (*i*.*e*. dynamic states) are shown on the right. The ATP binding sites are presented as molecular surfaces in ice blue. **h**, Representative western-blots of the expression levels of p-Erk1-2 and Erk1-2 in primary chondrocytes isolated from femoral cartilage of *Fgfr3*^*Y367C/*+^mice. **i**, Changes in the ratio between p-Erk1-2 and Erk1-2 (n=5). **j**, Representative western-blots of the expression levels of p-P38 and P38 in primary chondrocytes isolated from femoral cartilage of *Fgfr3*^*Y367C/*+^mice. **k**, Changes in the ratio between p-P38 and P38 (n=5). Data are quoted as the mean ± SEM. ns, not significant; *\*p<0*.*05*; *\*\*p<0*.*01*; *\*\*\*p<0*.*001*; *\*\*\*\*p<0*.*0001* in a two-tailed, unpaired t-test. The gray density analysis for the data of western-blot was done by using ImageJ software.

We evaluated the propensity of (-)-epicatechin to interact with FGFR3 downstream signaling pathways (ERK1, ERK2 and P38 kinases). We found that (-)-epicatechin remained bound to ERK1, ERK2 and P38 with only small structural variations from the starting dynamic state (static state) (Fig. 2f,g and Supplementary Fig. 4, Supplementary Table 1). To validate the interaction of (-)-epicatechin with FGFR3’s downstream signaling pathways in cartilage cells, we quantified the level of Erk1-2 and P38 phosphorylation in primary chondrocytes isolated from *Fgfr3*^*Y367C/*+^ mice^21^ treated with (-)-epicatechin. Compared with untreated *Fgfr3*^*Y367C/*+^ chondrocytes, the level of Erk1-2 phosphorylation (Fig. 2h, i) and P38 phosphorylation (Fig. 2j, k) were significantly lower in mutant chondrocytes treated with (-)-epicatechin. To definitively eliminate procyanidin C1 as drug candidate, *in silico* analyses demonstrated that procyanidin C1 failed to interact with Erk1, Erk2 and P38 (Supplementary Fig. 5 and Supplementary Table 2) and procyanidin C1 did not reduce the level of Erk1-2 phosphorylation in both primary *Fgfr3*^*Y367C/*+^ murine and human *FGFR3-Y373C* chondrocytes (data not shown). These findings indicate that the (-)-epicatechin might be a promising compound for repressing the constitutive activation of FGFR3’s downstream signaling pathways in cartilage cells.

### (-)-Epicatechin enhances bone growth and regulates ciliogenesis and growth plate organization

To assess the relevance of using polyphenol compound to treat cartilage tissue, we compared the growth of femurs isolated from dwarf *Fgfr3*^*Y367C/*+^ embryos (E16.5)^21^ in *ex vivo* cultures for six days in the presence of fraction 5 from *Theobroma cacao* (positive control), procyanidin C1 (used as a negative control), (-)-epicatechin and vehicle (Fig. 3a). The mean of growth is 1188 ± 33 µm for *Fgfr3*^+*/*+^ (Fig. 3b) and 423 ± 16 µm for *Fgfr3*^*Y367C/*+^ explants femurs at day 6 (Fig. 3c). The growth of (-)-epicatechin-treated *Fgfr3*^*Y367C/*+^ femurs was quite similar to fraction 5-treated *Fgfr3*^*Y367C/*+^ femurs (598 ± 35 µm) whereas procyanidin C1-treated *Fgfr3*^*Y367C/*+^ femurs (382 ± 19 µm) remains unchanged compared to *Fgfr3*^*Y367C/*+^ untreated femurs (Fig. 3c, d). (-)-Epicatechin treatment for 6 days has improved by 1.41-fold the growth of the mutant *Fgfr3*^*Y367C/*+^ femurs. At the end of the treatment, examination of the cartilage’s general layout and cell distribution revealed that the area of *Fgfr3*^*Y367C/*+^ femoral cartilage in treated distal and proximal femurs were larger (1.30.10^6^ ± 0.03.10^6^ µm^2^) than in non-treated femurs (1.12.10^6^ ± 0.05.10^6^ µm^2^) (Fig. 3e,f). Using frozen sections of *ex vivo* control and mutant femur growth plate, we examined the primary cilia’s length in the type-X-collagen-negative (proliferative) zone (Fig. 3g,h) and we confirmed that the primary cilium length was lower in the *Fgfr3*^*Y367C/*+^ proliferative zone (1.46 ± 0.01 µm) than in control *Fgfr3*^+*/*+^ (1.65 ± 0.01 µm) (Fig. 3g,h). Treatment with (-)-epicatechin corrected the primary cilium length in *Fgfr3*^*Y367C/*+^ chondrocytes (1.63 ± 0.01 µm) similarly to non-treated *Fgfr3*^+*/*+^ control chondrocytes (Fig. 3g,h). Measuring the x-y major axis of the cell’s longitudinal position and the angle ϕ with respect to the cell’s x-y axis, we observed that the angle ϕ was lower (27.1 ± 19.8°) in the *Fgfr3*^*Y367C/*+^ growth plate - suggesting that the primary cilium was unable to erect in the proliferative growth plate chondrocyte (Supplementary Fig. 6). (-)-Epicatechin restored the angle ϕ to values (39.7 ± 24.2°) close to those seen in *Fgfr3*^+*/*+^ experiments (41.6 ± 27.1°) (Supplementary Fig. 6).

**Figure 3.**
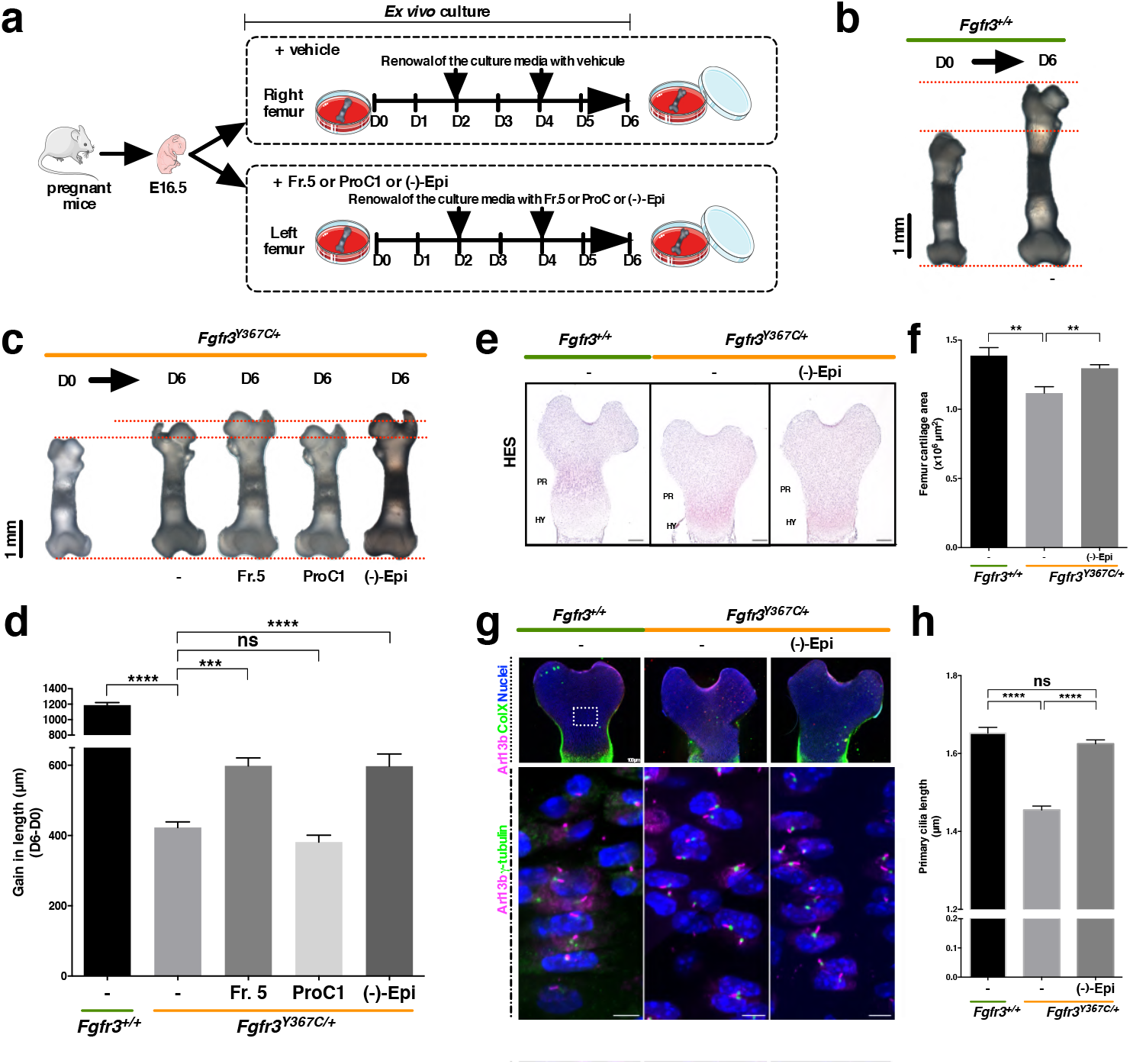
(-)-Epicatechin increases *ex vivo Fgfr3*^*Y367C/*+^ femur growth through primary cilia elongation. **a**, Sequential representation of *ex vivo* culture of femurs isolated from mouse embryos (E16.5). **b**, Representative images of E16.5 *Fgfr3*^+*/*+^ femurs in *ex vivo* culture for 0 (D0) and 6 (D6) days. **c**, Representative images of E16.5 *Fgfr3*^*Y367C/*+^ femurs in *ex vivo* culture for 0 (D0) and 6 (D6) days exposed (or not) to *Theobroma cacao* fraction 5, procyanidin C1 and (-)-epicatechin. **d**, Graphical representation of the gain in length of *ex vivo* E16.5 *Fgfr3*^+*/*+^ and *Fgfr3*^*Y367C/*+^ femurs after exposition (or not) to *Theobroma cacao* fraction 5, procyanidin C1 and (-)-epicatechin (n=15). **e**, Representative images of HES-stained immunostained embryonic distal femurs. Scale bars: 200 µm. PR: proliferative zone; HY: hypertrophic zone. **f**, Graphical representation of the femur growth plate area in E16.5 *Fgfr3*^+*/*+^ (n=5) and *Fgfr3*^*Y367C/*+^ femurs in the presence (n=8) or absence (n=7) of (-)-epicatechin. **g**, Representative confocal image of the primary cilium immunolabelled with with Arl13b (magenta), type X collagen or γ-tubulin (green), and DAPI (blue) localized in control and *Fgfr3*^*Y367C/*+^ femurs at day 6. **h**, Graphical representation of the length of the primary cilium in chondrocytes from femur cultures at day 6 of *Fgfr3*^+*/*+^ (n=917) and *Fgfr3*^*Y367C/*+^ exposed (n=1667) or not (n=1575) to (-)-epicatechin. Data are quoted as the mean ± SEM. ns, not significant; *\*p<0*.*05*; *\*\*p<0*.*01*; *\*\*\*p<0*.*001*; *\*\*\*\*p<0*.*0001* in a two-tailed, unpaired t-test.

### (-)-Epicatechin modulates cell proliferation and differentiation in ACH growth plate cartilage

To determine which cartilage markers proteins are targeted by (-)-epicatechin to promote femur elongation, we examined (-)-epicatechin’s putative action on chondrocyte differentiation (as characterized by changes in the cells’ morphology and collagen expression). The type X collagen expression, a marker of late-stage hypertrophic chondrocyte differentiation is decreased in *Fgfr3*^*Y367C/*+^ femurs^21^. Interestingly, we showed enlarged hypertrophic chondrocytes in the (-)-epicatechin treated growth plate at day 6 (Fig. 4a). In a quantitative immunolabeling analysis of collagen type X (Fig. 4b), we did not see a significant change in the overall size of the hypertrophic zone with (-)-epicatechin treatment (1.5.10^5^ ± 0.1.10^5^ µm^2^) (Fig. 4b). As shown in the high-magnification inset image of the type-X-collagen-positive zone (Fig. 4a, right panel), we found that the hypertrophic cells were 38.1% larger in (-)-epicatechin-treated *ex vivo Fgfr3*^*Y367C/*+^ femurs (214 ± 15 µm^2^) than in non-treated ones (155 ± 12 µm^2^) (Fig. 4c). We also noted that the number of cells per surface was double up in non-treated *Fgfr3*^*Y367C/*+^ growth plate cartilage (41.5 ± 0.9 chondrocytes per 10,000 µm^2^) compared to *Fgfr3*^+*/*+^ growth plate cartilage (16.8 ± 0.6 chondrocytes per 10,000 µm^2^). After 6 days of *ex vivo* culture, the number of cells per surface was significantly decreased in (-)-epicatechin-treated *Fgfr3*^*Y367C/*+^ femurs (35.4 ± 0.5 chondrocytes per 10,000 µm^2^; Fig. 4d) - suggesting that (-)-epicatechin treatment can, to some extent, partially rescue the defective, final differentiation of the collagen-type-X-positive cells in the *Fgfr3*^*Y367C/*+^ growth plate.

**Figure 4.**
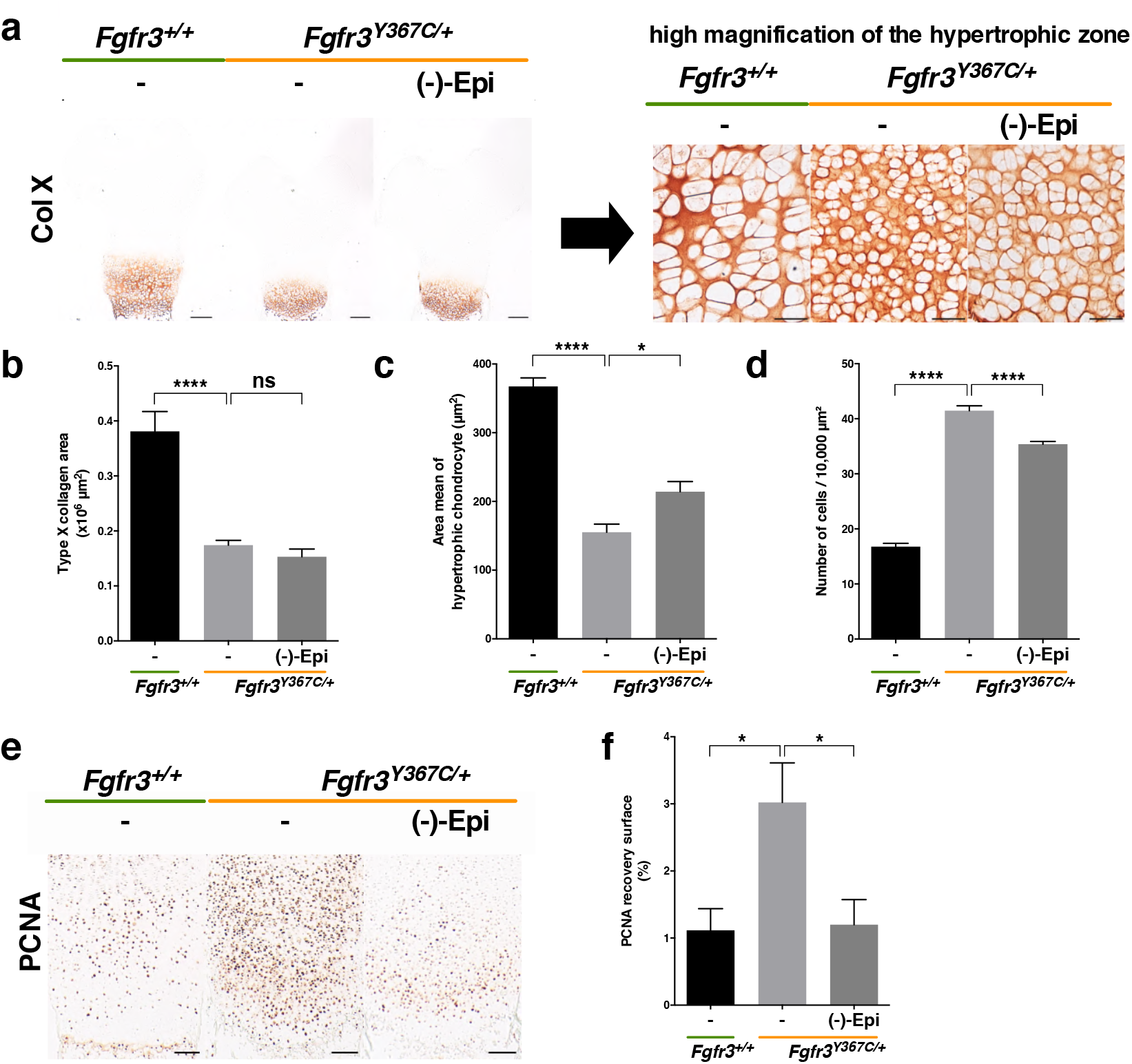
(-)-Epicatechin modulates hypertrophic and proliferative chondrocytes in *ex vivo Fgfr3*^*Y367C/*+^ femur. **a**, Representative images of type X collagen immunostaining of *Fgfr3*^+*/*+^ and *Fgfr3*^*Y367C/*+^ embryonic distal femurs (Scale bars: 200 µm). High-magnification inset of a representative image of type X collagen immunostaining in the hypertrophic distal zone of *Fgfr3*^+*/*+^ and *Fgfr3*^*Y367C/*+^ distal femurs (Scale bars: 50 µm). **b**, Graphical representation of type X collagen area immunostained E16.5 *Fgfr3*^+*/*+^ (n=5) and *Fgfr3*^*Y367C/*+^ distal femurs in the presence (n=8) or absence (n=7) of (-)-epicatechin. **c**, Graphical representation of the area of each hypertrophic chondrocyte of *Fgfr3*^+*/*+^ (n=5) and *Fgfr3*^*Y367C/*+^ distal femurs in the presence (n=5) or absence (n=5) of (-)-epicatechin. **d**, Graphical representation of the number of hypertrophic chondrocytes per surface of 10,000 µm2 of *Fgfr3*^+*/*+^ (n=80) and *Fgfr3*^*Y367C/*+^ distal femurs in the presence (n=80) or absence (n=5) of (-)-epicatechin. **e**, PCNA immunostaining of *Fgfr3*^+*/*+^ and *Fgfr3*^*Y367C/*+^ distal femurs. Scale bars: 100 µm. **f**, Graphical representation of the PCNA recovery surface of E16.5 *Fgfr3*^+*/*+^ (n=5) and *Fgfr3*^*Y367C/*+^ distal femurs in the presence (n=8) or absence (n=7) of (-)-epicatechin. Data are quoted as the mean ± SEM. ns, not significant; *\*p<0*.*05*; *\*\*p<0*.*01*; *\*\*\*p<0*.*001*; *\*\*\*\*p<0*.*0001* in a two-tailed, unpaired t-test.

We then hypothesized that (-)-epicatechin treatment mainly stimulated proliferative chondrocytes in the growth plate cartilage. We observed that proliferating cell nuclear antigen (PCNA) was overexpressed in the *Fgfr3*^*Y367C/*+^ cartilage growth plate (3.02 ± 0.59 % of PCNA recovery surface), relative to a control (1.12 ± 0.32 %) (Fig. 4e,f). Here, we found that PCNA expression levels were lower in (-)-epicatechin-treated *Fgfr3*^*Y367C/*+^ femurs (1.20 ± 0.37 % vs 3.02 ± 0.59 %) (Fig. 4f). Treatment with (-)-epicatechin for 6 days was seen to counter the abnormally high expression of PCNA - indicating that (-)-epicatechin normalizes the chondrocyte proliferation blockade in *Fgfr3*^*Y367C/*+^ femurs.

We also hypothesized that (-)-epicatechin treatment could stimulate chondrocytes differentiation in the growth plate cartilage. Chondrocyte differentiation is driven by many genes required to maintain healthy cartilage. The MAPK pathway (including P38 and Erk1-2) affects Sox9 transcription, chondrocyte differentiation, and cartilage matrix synthesis^29^. Sox9 as several transcription factors could also be regulated by P38, and the elevated level of phosphorylated P38 promote hypertrophic chondrocyte differentiation^30^. Here, we observed that abnormal activation of FGFR3 signaling promotes abnormal high Sox9 expression (as previously described^31,32^) and P38 and Erk1-2 phosphorylation levels in the growth plate cartilage of the femurs explants. Interestingly, treatment with (-)-epicatechin was associated with a lower level of Sox9, phosphorylated P38 and Erk1-2 in the *Fgfr3*^*Y367C/*+^ femur growth plate (Supplementary Fig. 7a,b, c). Overall, these proof-of-principle experiments showed that (-)-epicatechin modulates the activation of FGFR3’s major downstream signaling pathways.

Altogether, these data show that (-)-epicatechin has a beneficial effect on the proliferation/differentiation balance in *Fgfr3*^*Y367C/*+^ chondrocytes.

### (-)-Epicatechin modulates bone and primary cilia architecture in the *Fgfr3* mouse model of ACH

To confirm (-)-epicatechin’s *in vivo* effect on bone growth, we next sought to determine whether or not (-)-epicatechin could regulate bone growth in the *Fgfr3*^*Y367C/*+^ murine model of ACH^21^. The *Fgfr3*^*Y367C/*+^ mice were 1-day old upon treatment initiation. After administered daily, subcutaneous injections of (-)-epicatechin (0.1 mg.kg^-1^.d^-1^) or vehicle to *Fgfr3*^*Y367C/*+^ mice for 2 weeks (Fig. 5a), we X-rayed the animals (Fig. 5b). The mean naso-anal length was 4.9% greater in treated animals than in controls (38.51 ± 0.50 µm and 40.40 ± 0.46 µm for non-treated and (-)-epicatechin-treated *Fgfr3*^*Y367C/*+^ mice, respectively; Fig. 5c and Table 1). With regard to long bone growth, the femur and tibia increased by 6.82% (*p<0*.*0001*) and 6.08% (*p<0*.*001*) longer in (-)-epicatechin-treated *Fgfr3*^*Y367C/*+^ mice compared to controls, respectively (Fig. 5d,e and Table 1). Similarly, the lengths of the humerus, radius and ulna in treated *Fgfr3*^*Y367C/*+^ mice were respectively 3.21% (*p<0*.*01*), 5.09% (*p<0*.*01*) and 5.28% (*p<0*.*01*) longer than in non-treated controls (Fig. 5f-h and Table 1).

**Table 1.**
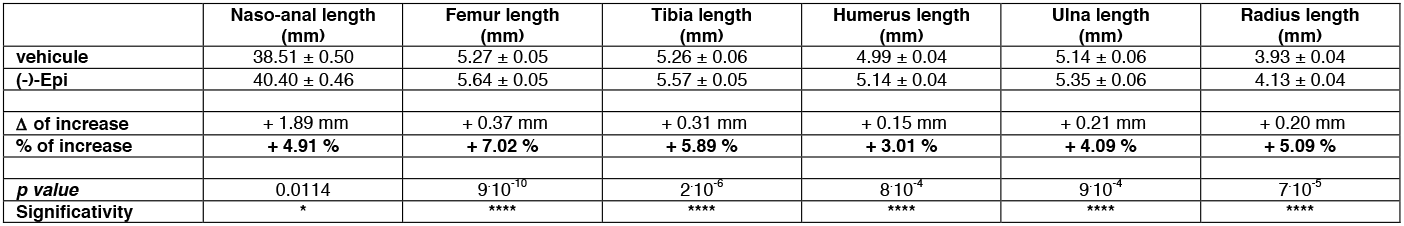
(-)-Epicatechin increases bone growth *in vivo*. The percentage growth gain of naso-anal, femur, tibia, humerus, ulna and radius lengths with detailed values for (-)-epicatechin-injected *Fgfr3*^*Y367C/*+^ animals. Data are quoted as the mean ± SEM. ns, not significant; *\*p<0*.*05*; *\*\*p<0*.*01*; *\*\*\*p<0*.*001*; *\*\*\*\*p<0*.*0001* in a two-tailed, unpaired t-test.

**Figure 5.**
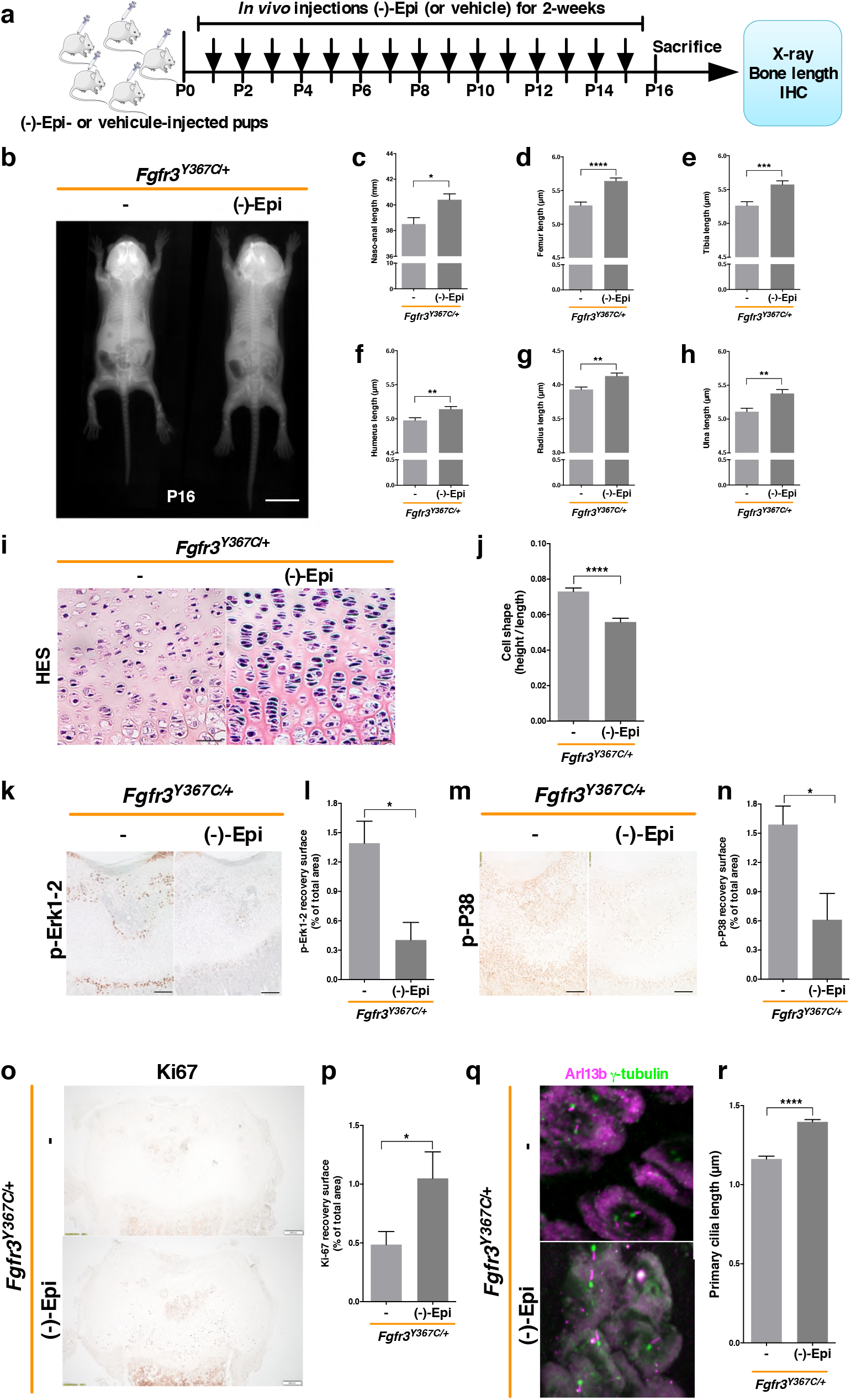
*In vivo*-injected (-)-epicatechin increases bone growth, modifies chondrocyte proliferation and differentiation and promotes primary cilium elongation. **a**, Sequential representation of *in vivo* protocol. **b**, Radiographs of *Fgfr3*^*Y367C/*+^ mice treated with vehicle or (-)-epicatechin for 16 days, and graphical representations of the naso-anal length (**c**), femur length (**d**), tibia length (**e**), humerus length (**f**), radius length (**g**) and ulna length (**h**). **i**, Representative images of HES staining of distal femurs. Scale bars: 50 µm. **j**, Graphical representation of the cell height/cell length ratio in the growth plate’s proliferative zone in *Fgfr3*^+*/*+^ (n=80) and *Fgfr3*^*Y367C/*+^ mice treated (n=80) or not (n=120) with (-)-epicatechin for 16 days. **k**, Representative image of p-Erk1-2 in *Fgfr3*^*Y367C/*+^ distal femurs in the presence (n=5) or absence (n=5) of (-)-epicatechin. Scale bars: 200 µm. **l**, Graphical representation of surface of p-Erk1-2 positive zone in *Fgfr3*^*Y367C/*+^ distal femurs in the presence (n=5) or absence (n=5) of (-)-epicatechin.. **m**, Representative image of p-P38 in *Fgfr3*^*Y367C/*+^ distal femurs in the presence (n=5) or absence (n=5) of (-)-epicatechin. Scale bars: 200 µm. **n**, Graphical representation of surface of p-P38 positive zone. **o**, Representative image of for Ki-67 in *Fgfr3*^*Y367C/*+^ distal femurs in the presence (n=5) or absence (n=5) of (-)-epicatechin. Scale bars: 200 µm. **p**, Graphical representation of surface of Ki-67 positive zone *Fgfr3*^*Y367C/*+^ distal femurs treated or not with (-)-epicatechin. **q**, Representative confocal microscopy image of primary chondrocytes from *in vivo Fgfr3*^*Y367C/*+^ femoral growth plate exposed or not to (-)-epicatechin. **r**, Graphical representation of the length of the primary cilium in chondrocytes after *in vivo* treatment (See also Supplementary Fig. 8 for enlarged images). Data are quoted as the mean ± SEM. ns, not significant; *\*p<0*.*05*; *\*\*p<0*.*01*; *\*\*\*p<0*.*001*; *\*\*\*\*p<0*.*0001* in a two-tailed, unpaired t-test.

The chondrocytes, from the *Fgfr3*^*Y367C/*+^ proliferative zone, were small, round and not structured into columns. (-)-Epicatechin’s *in vivo* mode of action is characterized by modification of the structural organization of the *Fgfr3*^*Y367C/*+^ growth plate cartilage, the chondrocytes were more aligned, flattened, and oriented in stacks (Fig. 5i). The *Fgfr3*^*Y367C/*+^ chondrocytes were significantly flatter after 2 weeks of treatment with (-)-epicatechin (ratio of 7.3.10^2^ ± 0.2.10^2^ compared to non-treated *Fgfr3*^*Y367C/*+^ mice: 5.6.10^2^ ± 0.2.10^2^; Fig. 5j). *In vivo* administration of (-)-epicatechin was able to normalize the expression of FGFR3 downstream effectors (Erk1-2, p38) (Fig. 5k-n) and proliferation (Ki67) (Fig. 5o,p).

*In vivo* (-)-epicatechin treatment has also a beneficial action of on primary cilia. We found that 2 weeks of (-)-epicatechin treatment was associated with markedly longer primary cilia in proliferative *Fgfr3*^*Y367C/*+^ chondrocytes (1.40 ± 0.01 µm) (Fig. 5q,r) and also an increase of ϕ angle values (39.7 ± 24.2°) (Supplementary Fig. 8).

These findings confirmed that systemically administered (-)-epicatechin *in vivo* penetrated the growth plate cartilage and strongly downregulated the expression of key regulators of chondrocyte proliferation and differentiation and modified the length of the primary cilia of chondrocytes *in vivo*. These results demonstrate that (-)-epicatechin can be used to control *in vivo* long bone elongation in FGFR3-related disease.

## Discussion

A growing body of research has identified a number of therapeutic approaches for the treatment of defective bone growth in achondroplasia, the most common form of dwarfism^2^. Our present results provide strong evidence for a potentially therapeutic effect of (-)-epicatechin on the cartilage growth plate and ciliogenesis in FGFR3-related disorders. Over the last few decades, the beneficial effects of polyphenols on human health have been increasingly well documented. Several preclinical studies have demonstrated that (-)-epicatechin is effective against sarcopenia^33^, alleviates inflammation in lipopolysaccharide-induced lung injury^34^, prevents the development of dilated cardiomyopathy^35^, and improves vascular function^36^. Here, we provide physiological evidence (in cell-based and murine models of ACH) and molecular evidence to show that (-)-epicatechin treatment modifies bone growth. Although fibroblast growth factor (FGF)’s role in skeletal development is quite well understood, little is known about the intracellular signaling that mediates the overactivation of FGFR3. Our data suggest that phosphorylated Erk1-2 and P38 and total Sox9 proteins are regulated in response to FGF signaling. Both Erk1-2 and P38 phosphorylation levels and the Sox9 expression level are elevated in the growth plate of *Fgfr3*^*Y367C/*+^ mice - indicating a major role for these proteins in *Fgfr3*-related skeletal dwarfism. The literature data show that P38, Sox9 and Erk1-2 upregulate hypertrophic differentiation^37,38^. Here, we found that *in silico* visualization of the FGFR3’s ATP-binding pocket showed clearly that (-)-epicatechin remained bound to ERK1, ERK2 and P38. Interestingly, *in vivo* (-)-epicatechin treatment decreased the activation of P38, Erk1-2 and Sox9 and was associated with a greater hypertrophic chondrocyte volume - known to be a major determinant of the longitudinal bone growth rate. We further showed that (-)-epicatechin interacts with the tyrosine kinase pocket of ERK1-2 and P38, which suggests the presence of a direct link to compound’s effects on the MAPK signaling pathway.

Our results also showed how (-)-epicatechin rescues proliferating columnar chondrocytes with a flattened, stacked appearance. The clonal expansion of the chondrocytes resulted in bone growth. At the cellular level, (-)-epicatechin rescued the defect in primary cilium elongation and thus modified cell division and columnar zone elongation. We therefore tested (-)-epicatechin as a treatment for directly affecting growth plate organization and proliferation by modifying the length and the ϕ angle of primary cilia in cartilage tissue and thus improving bone growth. We had previously demonstrated that sustained FGFR3 activity was associated with shorter cilia^6,8^, abnormal chondrocyte homeostasis, and disturbance of the growth plate’s columnar organization *via* chondrocyte rotation^8,9,11^. Our *in vitro* and *in vivo* results show that (-)-epicatechin was associated with longer chondrocyte primary cilia. Our morphometric studies of the growth plate showed that (-)-epicatechin treatment modified the chondrocytes’ alignment and the length and position of the primary cilia in the growth plate. These finding were supported by our transcriptomic analysis, highlighting (*i*) the abnormal expressions of several genes involved in ciliogenesis, the Ihh and PKA signaling pathways, and (*ii*) their normalization by (-)-epicatechin treatment. We were therefore not surprised to see that expression levels of genes linked to skeletal ciliopathies were abnormal in models of ACH; these genes included *Dync2li1* (the pathogenic gene in short rib polydactyly syndrome) and *Kif22* (the pathogenic gene in spondyloepimetaphyseal dysplasia with joint laxity)^39,40^.

Given the primary cilia’s essential role in Hh signaling and the known interactions between members of the FGF and Hh pathways^3^, it is likely that inhibition of the chondrocyte proliferation by upregulated FGF signaling is caused (at least in part) by the inactivation of Ihh signaling^41^. Our transcriptomic data confirmed that Hh signaling pathway is dysregulated in *Fgfr3*^*Y367C/*+^ mice. Ptch1 is essential for limb development^42,43^ and for interaction with Smoothened and Hh signaling through the Gli1/2/3 regulators^44–47^. This upregulation of *Ptch1* is in agreement with data from another mouse model of an *Fgfr3* gain-of-function mutation^48^. The present study demonstrated that (-)-epicatechin treatment controls Ihh-related primary cilia signaling pathway by acting on the Ihh/Ptch1/Gli1,2,3/Smo-cascade in cartilage. We also highlighted a putative role of the PKA pathway and *Adcy7* in *Fgfr3*^*Y367C/*+^ growth plate cartilage. Given that PKA was found to regulate Hh signaling in primary cilia and adenylate cyclase was clearly identified in skeletal primary cilium^49^, we hypothesize that PKA interferes with Ihh signaling in FGFR3-related disorders. Taken as a whole, our results demonstrate that (-)-epicatechin restores the impairment of the various genes controlling cilium elongation by acting on both the Ihh and PKA signaling pathways involved in bone growth regulation.

A better mechanistic understanding of the regulation of primary cilia formation *in vivo* provides the foundations for therapeutic intervention in FGFR3-related disorders. Although the current treatment for ACH restores chondrocyte differentiation *via* MAPK pathway, it fails to restore the columnar arrangement of the chondrocytes^13,14^; this contrasts with (-)-epicatechin, which (in addition to chondrocyte differentiation) restores the columnar arrangement by rescuing the defects in the primary cilium through regulation of primary cilia-related genes and thus enabling growth plate elongation. These data emphasize the relevance of (-)-epicatechin’s action in controlling both proliferation and differentiation. We consider that the modulation of FGFR3-activated signaling pathways by (-)-epicatechin provides a rationale for developing treatments for ACH.

## Materials and Methods

### Ethics statement

All the animal procedures and protocols were approved by the local animal care and use committee. Approval number APAFIS#24826-2018080216094268 v5, in compliance with EU directive 2010/63/EU for animals.

### Transcriptomics

Details of the *Fgfr3*^*Y367C/*+^ mouse models with C57BL/6 background have been described^21^. Three pairs of *Fgfr3*^*Y367C/*+^ and *Fgfr3*^+*/*+^ controls littermates were produced at four time points (d7, d14, d21 and d28). Mice genotypes were ascertained by PCR as described previously^21^. Mice femoral heads were collected at slaughter and frozen in liquid nitrogen. Samples were crushed, and RNA was extracted using a Qiagen RNeasy Mini kit and Qiagen’s animal tissue protocol. All RNA samples had an RNA integrity number above 8 and were analyzed on Affymetrix Mouse Genome 430 2.0 arrays by our service provider (PartnerChip, Evry, France). Processed and raw data were submitted to the Gene Expression Omnibus database (ID: GSE145821). Statistical analyses were performed using BioConductor2.5 and normalized using the gcrma package and were log2-transformed. Probesets differentially expressed by *Fgfr3*^*Y367C/*+^ *vs Fgfr3*^+*/*+^ animals during their development were identified by computing contrasts with a genotype*timepoint design. To define the gene expression profiles, we also computed interactions and linear, quadratic and cubic polynomial contrasts. P-values were adjusted for multiple testing using the eBayes function and the false discovery rate was computed using the decideTests function. Annotations were retrieved from the MGI database. Gene Ontology annotations were used to identify genes involved in cilium organization and functioning. Genes with at least 3 types of proof were also retrieved from Cildb^22^. Heat maps were generated online using Morpheus software.

### Chromatography and ESI-TOF mass spectrometry detection

The compounds in *Theobroma cacao* extract fractions were separated as described previously^16^, with one modification: the HPLC system was coupled to a TOF mass spectrometer equipped with an ESI interface, operating in negative ion mode with a capillary voltage of +3.5 kV. The optimum values of the other parameters were as follows: drying gas temperature, 200°C; drying gas flow, 10 L.min^-1^; and nebulizing gas pressure, 2.3 bar. The mass range for detection was 50–1200 *m/z*. To ensure repeatability, samples were injected in triplicate.

### NMR analysis

The sample was dissolved in DMSO-d6 and transferred into an oven-dried 5 mm NMR tube. The NMR spectra were recorded at 293 ± 0.1 K on a Bruker Avance III 600 spectrometer operating at a proton frequency of 600.13 MHz and fitted with a 5 mm QCI quadruple resonance pulse field gradient cryoprobe. The multiplicities observed were labeled as s = singlet; d = doublet; dd = doublet of doublets; t = triplet; m = multiplet; and bs = broad singlet. The sample was measured in 8 dummy scans prior to 128 scans. The acquisition parameters were as follows: size of free induction decay (FID) = 64K, spectral width = 20.5 ppm, acquisition time = 2.73 s, relaxation delay = 10 s, receiver gain = 20.2, and FID resolution = 0.25 Hz. A pre-saturation pulse sequence (Bruker 1D noesygppr1d) was used to suppress the residual H_2_O signal *via* irradiation at the H_2_O frequency during the recycle and mixing times. The resulting spectrum was automatically phased, baseline-corrected, and calibrated against the trimethylsilyl-2,2,3,3-tetradeuteropropionate signal at 0.0 ppm. The t1 time was set to 4 µs, and the mixing time (d8) was set to 10 ms. The spectrometer transmitter was locked to the DMSO-d6 frequency. Spectra were processed using TopSpin™ software (version 3.1, Bruker, Germany). 1H–1H total correlation spectroscopy (TOCSY) spectra, 1H-13C heteronuclear single quantum coherence (HSQC) spectra, and 1H-13C heteronuclear multiple bond coherence (HMBC) spectra were recorded using standard Bruker sequences. The TOCSY spectrum was obtained by applying a relaxation delay of 2.0 s, a spectral width in both dimensions of 7194.25 Hz, and a receiver gain of 64.0. It was then processed using a sine-bell window function (shifted sine bell = 2.0). The HSQC spectrum was acquired using a relaxation delay of 1.0 s, and a spectral width of 7211.54 Hz in F2 and 24900.71 Hz in F1. A quadratic sine window function (shifted sine bell = 2.0) was used for the HSQC spectrum. The HMBC spectrum was recorded with the same parameters as for the HSQC spectra, except that the spectral width in F1 was 37729.71 Hz. The coupling constant for HSQC experiments was set to 145 Hz, and 145 and 8 Hz (long range) for the HMBC experiments.

### Primary and immortalized human chondrocytes and mouse chondrocyte cultures

Primary and immortalized mutant cells were obtained and cultured as described previously^8,20^. Human and mouse cells were incubated with FGF2 (100 ng.mL^-1^) for 5 min, and then with *Theobroma cacao* extract fraction 5 (100 µg.mL^-1^) or (-)-epicatechin (80 µg.mL^-1^) (Sigma-Aldrich, HW101708-1) for 30 min. Cultured chondrocytes were immunolabelled as previously described^8^.

### Total RNA extraction and RT-qPCR

A reverse transcriptase quantitative polymerase chain reaction (RT-qPCR) was used to evaluate RNA expression after (-)-epicatechin treatment. RNA was extracted using the RNeasy Mini Kit (Qiagen) according to the manufacturer’s instructions. Measured quantity of RNA (500 ng) was reverse-transcribed using SuperScript™ II Reverse Transcriptase (Invitrogen). RT-qPCR was performed with a ViiA-7 system (Applied Biosystems) using SYBR Green (Life Technologies) for fluorescence detection. Primers were designed using the Primer3Plus website. RT-qPCR data were analyzed using the 2^-ΔΔCT^ method and β-actin was used as the housekeeping control. The primer sequences used were as follows:

**Dync2li1-Fwd:**^5’^AGTCAGCAAAGCCGACCTTAGCG^3’^;

**Dync2li1-Rev:** ^5’^TCCAGCAAGGAGGTTCCTCCACC^3’^;

**Kif24-Fwd:** ^5’^TCGCCAACATCTCCCCAAGCC^3’^;

**Kif24-Rev:** ^5’^TGGTAGCTGAAGCACAACACTTAACGC^3’^;

**Kif22-Fwd:** ^5’^AGGGAAGCCAAGGGACCCCC^3’^;

**Kif22-Rev:** ^5’^GTGCCTCCTGGGCCAACACC^3’^;

**Ptch1-Fwd:** ^5’^CTGGCAGCCGAGACAAGCCC^3’^;

**Ptch1-Rev:** ^5’^CCGGTGAGGCCGGATGTTGG^3’^;

**Ihh-Fwd:** ^5’^GGCGCGCTTAGCAGTGGAGG^3’^;

**Ihh-Rev:** ^5’^CTGGGCTCCGGCAGGAAAGC^3’^;

**Smo-Fwd:** ^5’^CTGGACCAAGGCCACCCTGC^3’^;

**Smo-Rev:** ^5’^TTCTGCAGCAGCTCACGCCG^3’^;

**Gli1-Fwd:** ^5’^CGCGCCAAGCACCAGAATCG^3’^;

**Gli1-Rev:** ^5’^TGTTTGCGGAGCGAGCTGGG^3’^;

**Gli2-Fwd:** ^5’^CGAGACCAACTGCCACTGGGC^3’^;

**Gli2-Rev:** ^5’^TGGGCCTTGAAGGGCTTCTGC^3’^;

**Gli3-Fwd:** ^5’^CGCTCATTGCACAGCAGCCC^3’^;

**Gli3-Rev:** ^5’^GGATGGCCCCTGCACCAGC^3’^;

**Atp8a2-Fwd:** ^5’^TGGACCTGGCGCTCTCATGC^3’^;

**Atp8a2-Rev:** ^5’^CCGATGGCCAGGGTGATGGC^3’^;

**Prkar2b-Fwd:** ^5’^ACCTCTCCTGGTGCTCTGTGGG^3’^;

**Prkar2b-Rev:** ^5’^CTTGAGGAACGGCAGGGACTCG^3’^;

**Adcy7-Fwd:** ^5’^GGCCTTGGTGGCCTACCTGC^3’^;

**Adcy7-Rev:** ^5’^CGGAGCAAGGGTGTGGCTGG^3’^;

**β-Actin-Fwd:** ^5’^CATGTTTGAGACCTTCAACAC^3’^;

**β-Actin-Rev:** ^5’^GCCATCTCCTGCTCGAAGTCTAG^3’^.

### Transient transfections with cDNA FGFR3 constructs

HEK293 cells at 70%–80% confluence were transiently transfected with FGFR3 human constructs (FGFR3-Y373C, FGFR3-G380R, FGFR3-K650E). Transfected cells were incubated with (-)-epicatechin overnight, and then lysed in RIPA buffer (50 mM Tris-HCl pH 7.6, 150 mM NaCl, 0.5% NP40, and 0.25% sodium deoxycholate, supplemented with protease and phosphatase inhibitors; Roche). Proteins were immunoprecipitated by the addition of 3 µl of rabbit anti-FGFR3 (Sigma-Aldrich, F0425)/500 µg protein with protein A-agarose (Roche). Fished-out FGFR3 protein was eluted and then denatured at 95°C for 10 minutes in NuPAGE LDS sample buffer (Life Technologies, NP0008) and β-mercaptoethanol 2.5%.

### Immunoblot analysis

Whole-cell lysates and immunoprecipitated proteins were subjected to NuPAGE 4%-12% bistris acrylamide gel electrophoresis (Life Technologies). Using standard protocols, we probed the blots with the following primary antibodies: anti-phosphotyrosine (Cell Signaling Technology), anti-phosphorylated Erk1-2 (Thr180/Tyr182) (Cell Signaling Technology, #4370), anti-total Erk1-2 (Sigma-Aldrich, M5670), anti-phosphorylated p38 (Thr180/Tyr182) (Cell Signaling Technology, #9205), anti-total p38 (Cell Signaling Technology, #2708), anti-total PCNA (Abcam, ab29-100) and anti β-actin (Millipore, MAB1501). Proteins were immunodetected with an anti-rabbit IRDye^®^ 800CW antibody (LI-COR Biosciences, 926-32211) and an anti-mouse IRDye^®^ 680RD antibody (LI-COR Biosciences, 926-68070). Immunoblots were quantified with ImageStudioLite software (LI-COR Biosciences).

### Computational details

The starting coordinates of ERK1, ERK2 and P38 were downloaded from the Protein Data Bank with respective accession codes: 4QTB, 5NGU and 4L8M. Ligand (-)-epicatechin was built using Maestro software and optimized with Amber software. Molecular docking calculations were performed with Autodock 4.2 software. We selected the docking pose that presented the lowest free energy of binding in the most populated cluster. The stability of the complexes formed by (-)-epicatechin and ERK1, ERK2, P38 and FGFR3 kinases were further explored through molecular dynamics simulations, using Amber software and the FF14SB/gaff2 force-field. For each system, the simulation started with an energy optimization of the water solvent and was followed by an overall minimization and a heating phase (from 0 to 300 K). Next, a molecular dynamics production was performed with the NTP ensemble, leading to a 50 ns trajectory. After the 50 ns period of conventional molecular dynamics, all systems underwent an additional 50 ns gaussian accelerated molecular dynamics protocol.

### *Ex vivo* femur culture system

Femurs were cultured *ex vivo*, as described previously^20^. The left femur was cultured in the presence of *Theobroma cacao* extract fraction 5 (10 µg.mL^-1^) or procyanidin C1 (10 µg.mL^-1^) or (-)-epicatechin (10 µg.mL^-1^), and compared with the non-treated right femur. The bone’s length was measured on day 0 (D0) and day 6 (D6).

### Histological and immunohistochemical analyses of paraffin sections

Femur explants were fixed in 4% paraformaldehyde and embedded in paraffin. Serial 5 µm sections were stained with hematoxylin-eosin-safran reagent, using standard protocols. For immunohistochemical assessment, sections were labeled with the following antibodies and a Dako Envision Kit: anti-Col X (BIOCYC, N.2031501005; 1:50 dilution), anti-Sox9 (polyclonal antibody, Santa Cruz Biotechnology Inc., catalog D0609; dilution 1:75), anti-phosphorylated Erk1-2 (Thr180/Tyr182) (Cell Signaling Technology, #4370; 1:100 dilution), anti-phosphorylated P38 (Abcam, Ab4822; 1:200 dilution) and anti-Ki67 (Abcam, Ab16667; 1:3,000 dilution). Images were captured with an Olympus PD70-IX2-UCB microscope and quantified using cellSens software.

### Histological and immunohistochemical analyses of frozen sections

Femur explants or femurs isolated from mice at P16 were fixed in methanol chilled at -20°C for 5h or 24h, respectively. After incubation in EDTA 0.5 M, pH 8 for 72 h or 2-weeks, femurs were placed in sucrose 30% for 24 h, transferred into OCT compound at room temperature, and frozen in isopentane at -45°C. The 50 µm tissue sections were permeabilized with Triton X-100 0.3% for 30 min, and immunolabeled with rabbit IgG anti-Arl13b (Proteintech #17711-1-AP, IF 1:100) or mouse IgG_1_ anti-γ-tubulin (Sigma-Aldrich #T6557, 1:100) primary antibodies. The primary antibodies were detected with goat anti-mouse IgG_1_ coupled to AlexaFluor 488 (LifeTechnologies, 1:400) and anti-rabbit IgG coupled to AlexaFluor 647 (LifeTechnologies, 1:400). Tissue sections were mounted with DAPI-Fluoromount G^®^ (CliniSciences). Three-dimensional images of the growth plate were obtained using a spinning disc confocal microscope. Images were displayed using FIJI and the FigureJ plugin.

### The mouse model and drug treatments

The *Fgfr3*^*Y367C/*+^ mouse model with C57BL/6 background has been described previously^20^. Cartilage and bone analyses were performed on 16-day-old mice. The *Fgfr3*^*Y367C/*+^ mice were 1-day old upon treatment initiation, and received daily subcutaneous administrations of (-)-epicatechin (Sigma-Aldrich, HW101708-1) (0.1 mg.kg^-1^ body weight) or vehicle (3.5 mM HCl, 0.1% DMSO) for 2-weeks. Long bones were measured using a caliper (VWRi819-0013, VWR International).

### Statistical analysis

Differences between experimental groups were assessed in an analysis of variance (ANOVA) with Tukey’s post hoc test or a Mann–Whitney U test. The threshold for statistical significance was set to *p* ≤ 0.05. Statistical analyses were performed using GraphPad PRISM software. A paired Student’s t-test was used to compare two treatments in the same cell population. An unpaired-Student’s t-test was used to compare groups of mice or different primary chondrocyte preparations.

## Acknowledgements

We thank the Imagine Institute’s imaging facility and the SFR’s histology facility for their help with this work. This program received a state subsidy managed by the National Research Agency under the “Investments for the Future” program bearing the reference ANR-10-IAHU-01.

## Author contributions

LM, NK, CBL, LS and LLM designed the research. LM, NK, CBL, SD, LS, VE, MC, LL performed the *in vitro* and *ex vivo* experiments. NK, LS and MM performed the *in vivo* experiments. MCD, SFA and ASC designed the *Theobroma cacao* extraction. FB performed the *in silico* analyses. LM, NK, CBL prepared the figures. LM and LLM wrote the manuscript.

## Supplementary Figures and Supplementary Figures legends

**Supplementary Figure 1.**
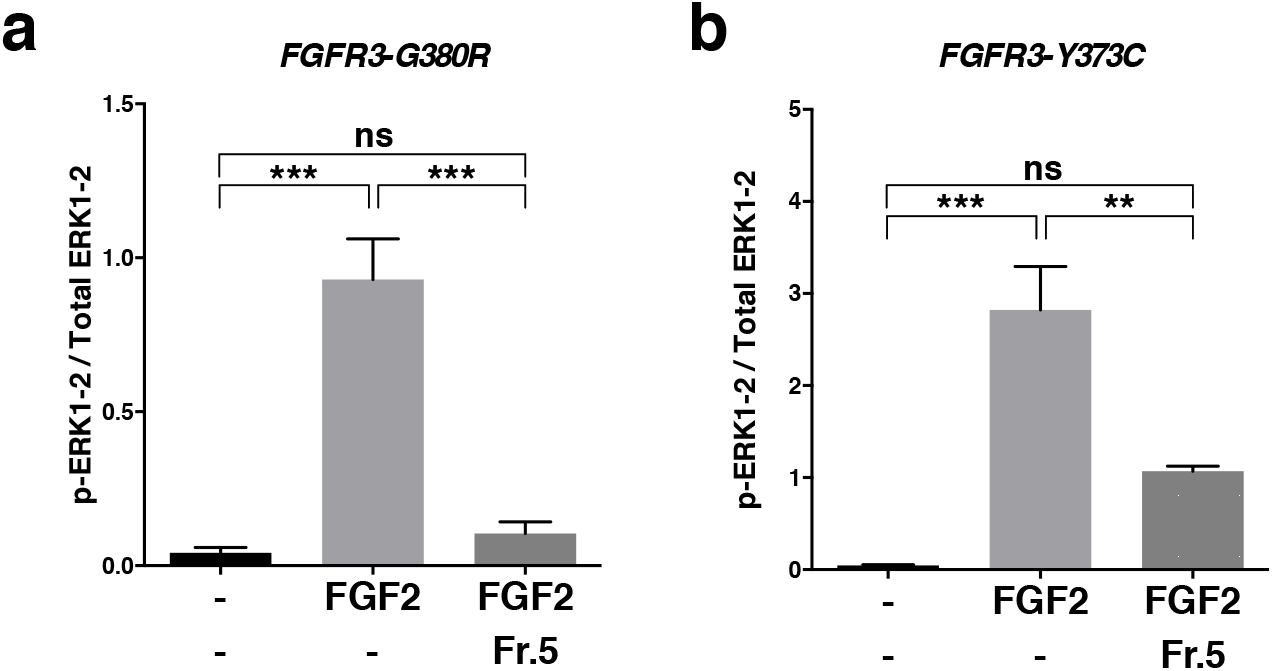
*Theobroma cacao* extract fraction 5 inhibits ERK1-2 phosphorylation in human chondrocytes. **a**, Graphical representation of the amounts of p-ERK1-2 and total ERK1-2 in human chondrocyte expressing ACH mutation (G380R) stimulated by FGF2 and/or *Theobroma cacao* extract fraction 5. **b**, Graphical representation of the ratio between p-ERK1-2 and total ERK1-2 in human chondrocyte expressing TD mutation (Y373C) stimulated by FGF2 and/or *Theobroma cacao* extract fraction 5. Data are quoted as the mean ± SEM. ns, not significant; *\*p<0*.*05*; *\*\*p<0*.*01*; *\*\*\*p<0*.*001*; *\*\*\*\*p<0*.*0001* in a two-tailed, unpaired t-test.

**Supplementary Figure 2.**
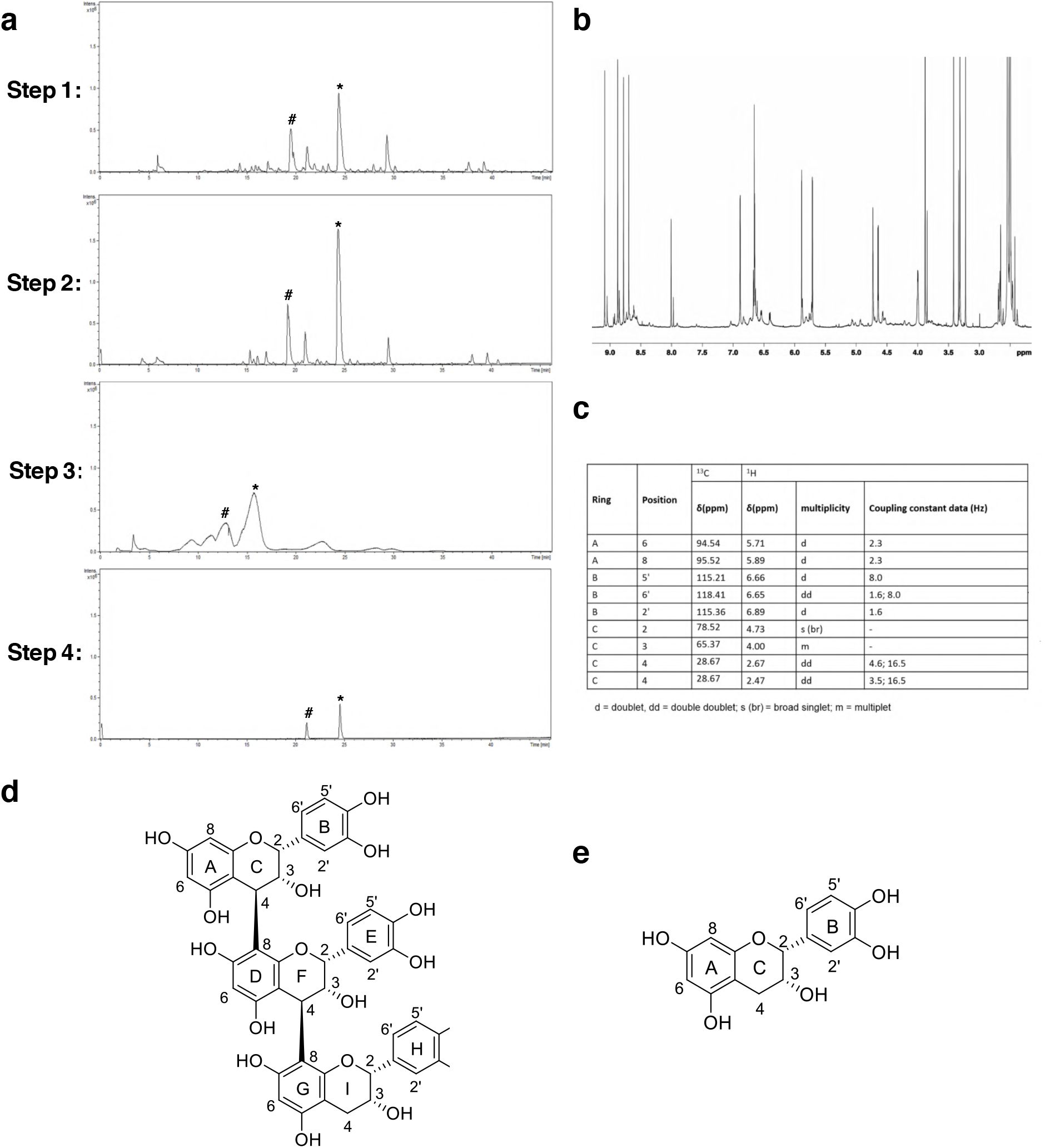
Purification and identification of procyanidin C1 and (-)-epicatechin from *Theobroma cacao* extract. **a**, Steps in the purification of (-)-epicatechin from *Theobroma cacao* extract (Step 1: an LC-MS base peak chromatogram (BPC) of *Theobroma cacao* DMSO extract. Step 2: BPCs of selected fractions from solid phase extraction (SPE) isolation. Step 3: BPC of HPLC semi-preparative purification of the SPE fraction. Step 4: BPCs of selected fractions purified using semi-preparative HPLC (Fraction (Fr) 5)); **b**, NMR ^1^H spectra of fraction 5 in DMSO-d_6_; **c**, ^1^H and ^13^C NMR chemical shift data for (-)-epicatechin from fraction 5 in d_6_-DMSO; **d**, The chemical structure of procyanidin C1, as identified in *Theobroma cacao* extract fraction 5 and **e**, The chemical structure of (-)-epicatechin, as identified in *Theobroma cacao* extract fraction 5. #: peak associated with procyanidin C1 from *Theobroma cacao* extract fraction 5; *: peak associated with (-)-epicatechin from *Theobroma cacao* extract fraction 5.

**Supplementary Figure 3.**
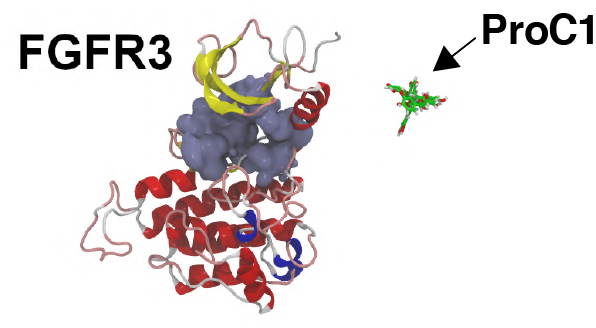
Representative structure of absence of complex between procyanidin C1 with FGFR3 kinase. The absence of complexe obtained after the molecular dynamics and hyperdynamics simulation stages (*i*.*e*. dynamic states). The ATP binding site is presented as molecular surfaces in ice blue.

**Supplementary Figure 4.**
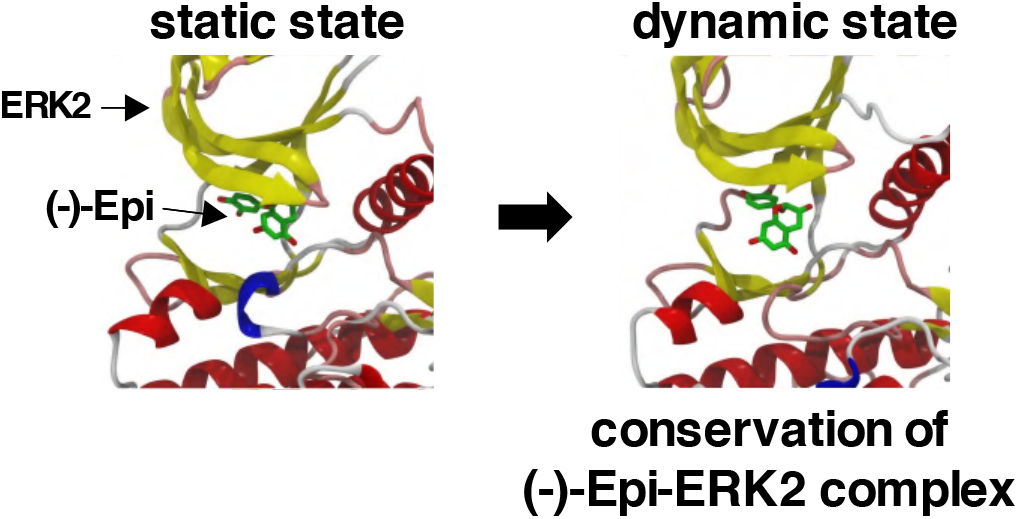
Representative structures of (-)-epicatechin complexed with ERK2 kinases. The complexes obtained after the molecular docking computations (*i*.*e*. static states) are shown on the left, and those obtained after the molecular dynamics and hyperdynamics simulation stages (*i*.*e*. dynamic states) are shown on the right. The ATP binding sites are presented as molecular surfaces in ice blue.

**Supplementary Figure 5.**
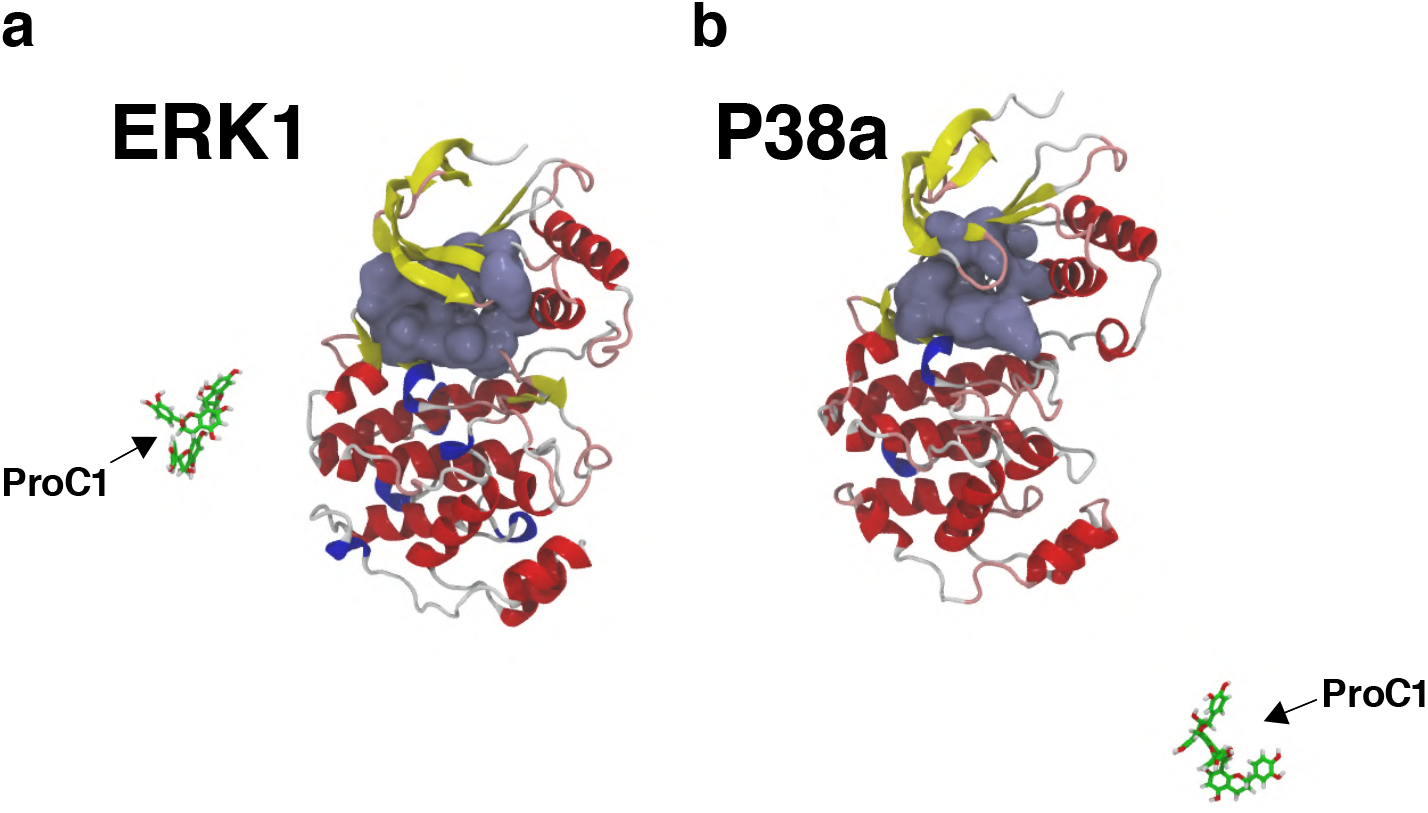
Representative structures of absence of complex between procyanidin C1 with ERK1 and P38 kinases. The absence of complexe obtained after the molecular dynamics and hyperdynamics simulation stages (*i*.*e*. dynamic states) are shown for ERK1 (**a**) and P38 (**b**) kinases. The ATP binding sites are presented as molecular surfaces in ice blue.

**Supplementary Figure 6.**
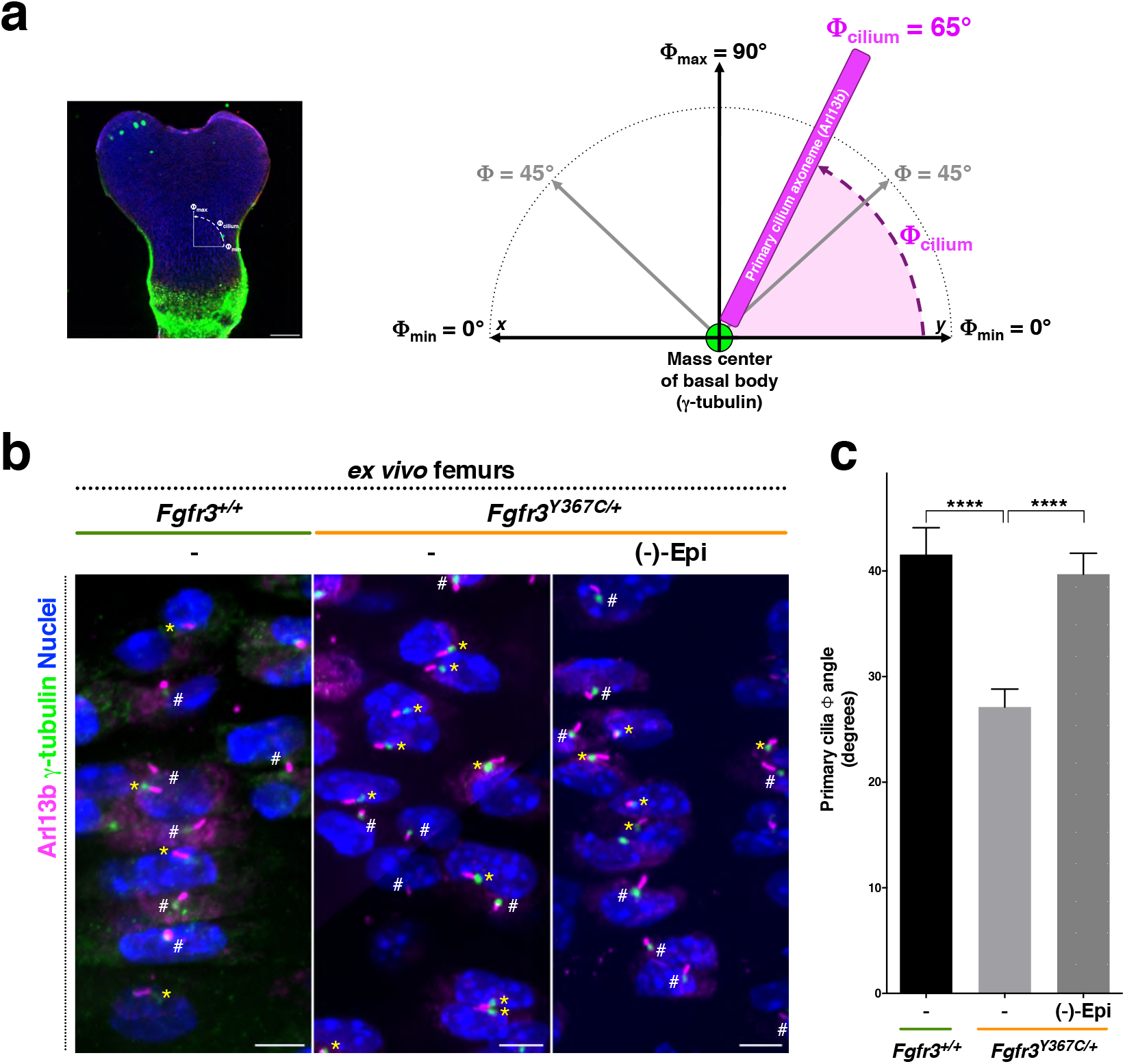
(-)-Epicatechin modifies primary cilium orientation angle defects *ex vivo*. **a**, Representive schema of the ϕ angle measurement. **b**, Representative confocal microscopy image of Arl13b (magenta), γ-tubulin (green), and DAPI (blue) in the primary cilium in E16.5 chondrocytes. Scale bars: 1 µm. White sharps: primary cilia with a random orientation (angle ϕ); yellow asterisks: primary cilia with a relatively low angle ϕ (< 20°) with bone growth elongation. **c**, Graphical representation of the orientation angle f for the primary cilium in chondrocytes from *ex vivo* femur cultures of *Fgfr3*^+*/*+^ (n=114) and *Fgfr3*^*Y367C/*+^ exposed (n=134) or not (n=150) to (-)-epicatechin. Data are quoted as the mean ± SEM. ns, not significant; *\*p<0*.*05*; *\*\*p<0*.*01*; *\*\*\*p<0*.*001*; *\*\*\*\*p<0*.*0001* in a two-tailed, unpaired t-test.

**Supplementary Figure 7.**
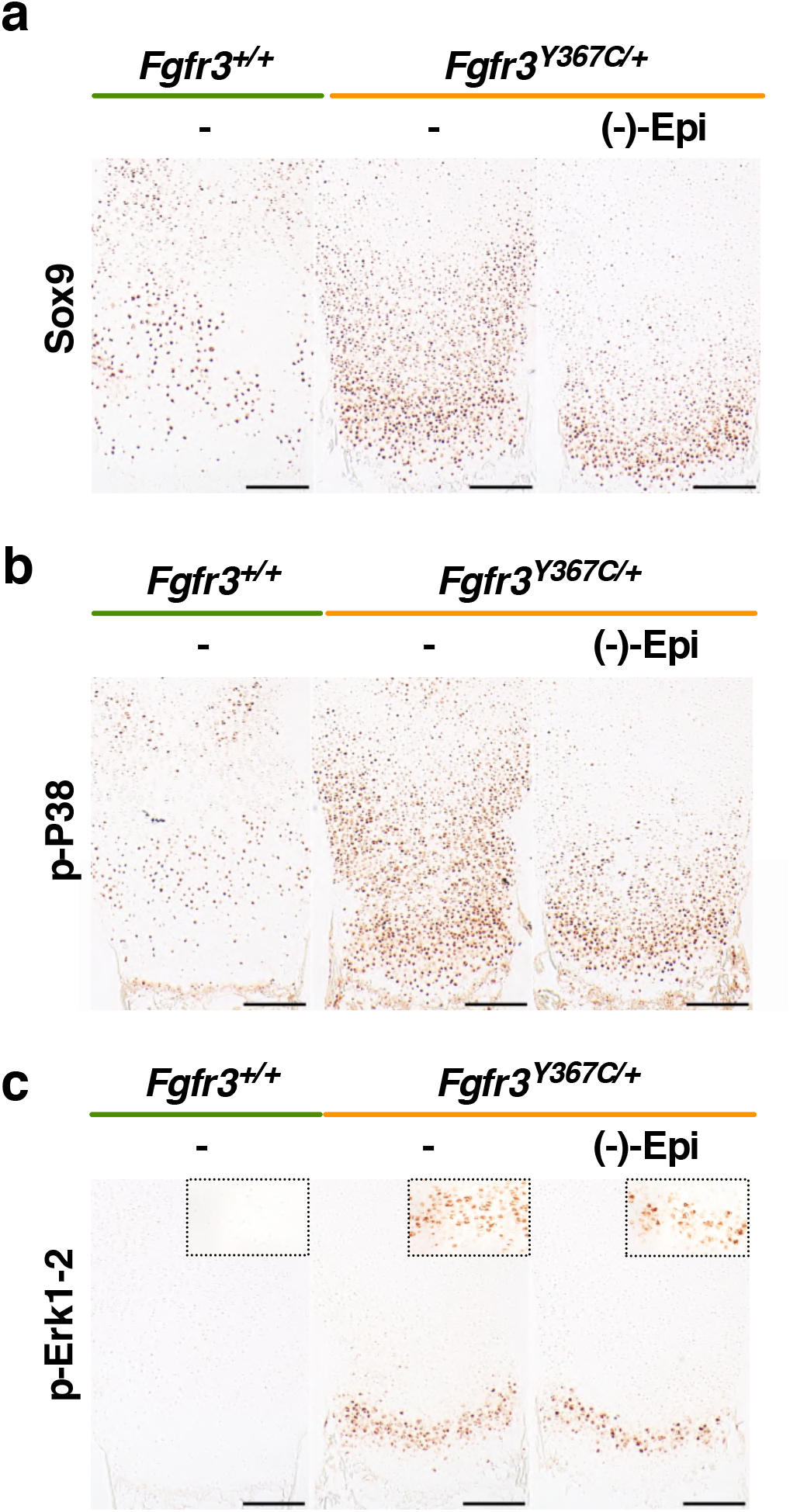
(-)-Epicatechin modifies the growth plate of distal *ex vivo Fgfr3*^*Y367C/*+^ femurs. **a**, Immunostaining for Sox9 on E16.5 *Fgfr3*^+*/*+^ and *Fgfr3*^*Y367C/*+^ distal femurs exposed or not to (-)-epictechin. **b**, Immunostaining for p-p38 on E16.5 *Fgfr3*^+*/*+^ and *Fgfr3*^*Y367C/*+^ distal femurs exposed or not to (-)-epictechin. **c**, Immunostaining for p-Erk1-2 (high magnification of the hypertrophic zone in the right top corner) on E16.5 *Fgfr3*^+*/*+^ and *Fgfr3*^*Y367C/*+^ distal femurs exposed or not to (-)-epictechin. PR: Proliferative zone. HY: Hypertrophic zone. Scale bars: 100 µm, 50 µm.

**Supplementary Figure 8.**
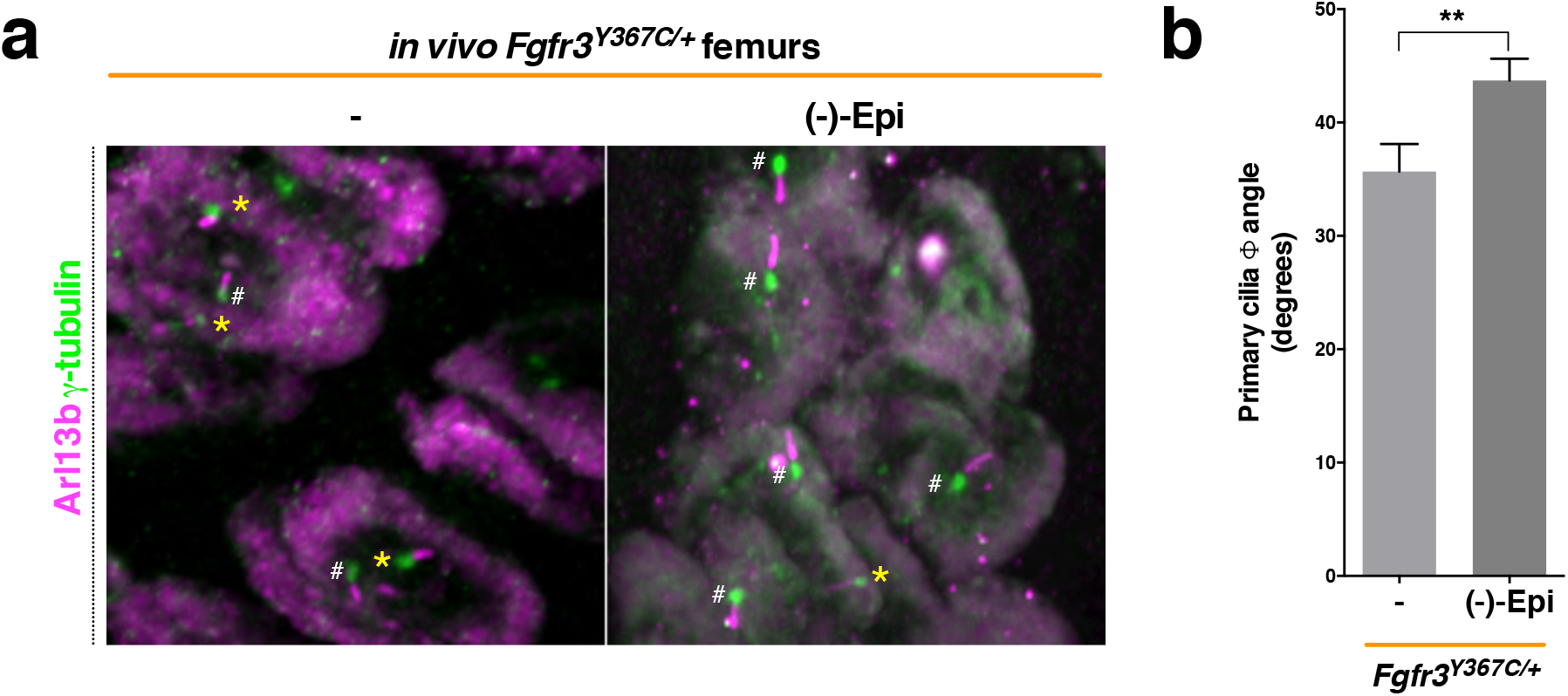
(-)-Epicatechin modifies also primary cilium orientation angle defects *in vivo*. **a**, Representative confocal microscopy image of primary chondrocytes from *in vivo Fgfr3*^*Y367C/*+^ mice immunolabelled by Arl13b (magenta), γ-tubulin (green), and DAPI (blue). White sharps: primary cilia with a random orientation (angle ϕ); yellow asterisks: primary cilia with a relatively low angle ϕ (< 20°) with bone growth elongation. **b**, Graphical representation of the primary cilium orientation angle ϕ in chondrocytes after *in vivo* treatment. Data are quoted as the mean ± SEM. ns, not significant; *\*p<0*.*05*; *\*\*p<0*.*01*; *\*\*\*p<0*.*001*; *\*\*\*\*p<0*.*0001* in a two-tailed, unpaired t-test.

**Supplementary Table 1.**
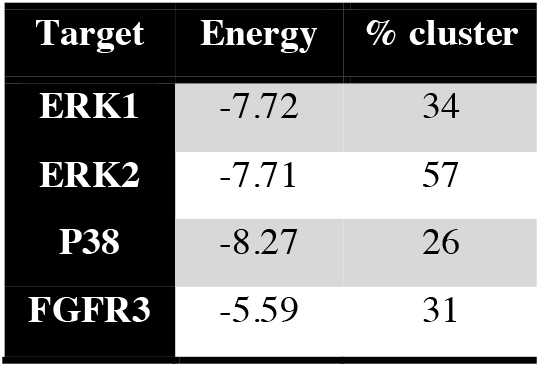
Docking binding energies (in kcal.mol^-1^) along with the percentage of cluster for (-)-epicatechin and the ERK1, ERK2, p38 and FGFR3 kinases.

| Target | Energy | % cluster |
| --- | --- | --- |
| ERK1 | -7.72 | 34 |
| ERK2 | -7.71 | 57 |
| P38 | -8.27 | 26 |
| FGFR3 | -5.59 | 31 |

**Supplementary Table 2.** Docking binding energies (in kcal.mol^-1^) along with the percentage of cluster for procyanidin C1 and the FGFR3, ERK1, ERK2 and P38 kinases.

| Target | Energy | % cluster |
| --- | --- | --- |
| FGFR3 | -5.7 | 9 |
| ERK1 | -5.1 | 6 |
| ERK2 | -2.8 | 16 |
| P38 | -7.5 | 7 |

**Supplementary Movie 1**. Molecular simulation of (-)-epicatechin from FGFR3 kinase.

**Gene Expression Omnibus, database ID: GSE145821**

